# Tag-stabilized fluorescent protein binders for endogenous cell type-specific protein labelling

**DOI:** 10.64898/2026.09.29.754891

**Authors:** ST Schnider, C Kring, L Maggi, MA Vigano, M Affolter, G Aguilar, A Kempf

## Abstract

Protein labelling using genetically encoded fluorescent protein fusions has revolutionized the study of protein localization and function. However, visualizing endogenous proteins in specific cell types remains challenging, as existing approaches often require overexpression or laborious genetic engineering. To address these limitations, we developed a library of fluorescent protein binders using existing nanobodies and single-chain variable fragments that recognize fluorescent proteins and short peptide tags, allowing for modular, cell type-specific labelling of genetically tagged endogenous proteins. We found that fluorescent protein binders generate background fluorescence in the absence of the target protein, reducing signal specificity. To overcome this limitation, we generated destabilized fluorescent protein binders directed against GFP, ALFA, GCN4 peptide and V5. These binders display increased antigen-dependent stability and reduced background fluorescence, thereby improving the signal-to-noise ratio of endogenous protein visualization. We demonstrate the detection of endogenous proteins in drosophila larvae and embryos, zebrafish embryos, as well as in specific neuronal populations of the adult drosophila brain. In the brain, tissue-specific labelling revealed previously unknown protein expression and subcellular localization in sleep-regulatory neurons. Together, our panel of tag-stabilized fluorescent protein binders provides a versatile, ready-to-use toolkit for background-free, cell type-specific visualization of endogenous proteins.

## Introduction

The analysis of protein localization and function is essential for elucidating the cellular mechanisms underlying development and organismal physiology. Fluorescence microscopy of proteins fused to fluorescent proteins has profoundly advanced the study of protein localization and dynamics (*1*). However, resolving the subcellular distribution of broadly expressed fluorescent protein (FP) fusions within intact tissues remains challenging because widespread fluorescence can obscure protein localization in individual cell types. This limitation can be partially overcome by restricted protein labelling, which enables the selective examination of the localization of broadly expressed proteins within defined cell populations (*2*). For example, in drosophila, cell type-specific protein labelling is commonly achieved by ectopically expressing FP fusions under the control of cell type-specific enhancers (*3*). Although these approaches enable selective visualization of proteins in defined cells (*4*), they rely on ectopic expression, which often results in protein levels that differ from endogenous abundance and can perturb protein localization or function (*5*). More recently, gene-editing technologies have enabled cell type-specific expression of endogenously tagged proteins (*6–9*). While these approaches have substantially improved the analysis of endogenous protein localization, they require extensive locus-specific engineering, limiting their routine application.

Protein binders, including single-domain antibodies (nanobodies), single-chain variable fragments (scFvs) and monobodies (FingRs), among others, have emerged as powerful tools to monitor and manipulate proteins (*10*). These binders can recognize epitopes with high affinity and selectivity and can themselves be fused to fluorescent proteins to indirectly label target proteins (*11–16*). Genetically encoded fluorescent binders have been applied for cell type-specific protein visualization in mouse (*13, 17*), zebrafish (*18*) and drosophila (*9, 19–22*). Binders directed against endogenous epitopes eliminate the need for genetic tagging altogether (*13, 18, 23*). However, because each target requires the identification and careful characterization of a dedicated binder, this strategy remains difficult to generalize.

A major advance in intracellular protein visualization was the development of the GFP chromobody, a GFP-directed nanobody fused to a FP that enables the detection of GFP-tagged proteins (*12*). The subsequent isolation of binders recognizing short peptide epitopes further expanded the versatility of this approach (*22, 24–30*). More recently, epitope tag-binder systems have been adapted for the visualization of endogenous proteins in multicellular organisms following minimal genetic modification of the target protein with a single epitope tag (*9, 21, 22, 31–33*). In drosophila, the high-affinity ALFA nanobody enables the visualization of the relatively abundant protein Myosin II (*9*). However, reliable visualization of low-abundance proteins, such as the synaptic protein Neurexin-1, remains considerably more challenging (*21*).

A major limitation in the visualization of endogenous proteins is the intracellular stability of some fluorescent protein binders. Stable fluorescent binders can accumulate even in the absence of their target, thereby generating background fluorescence that can obscure target-associated signals (*9, 21, 34, 35*). Conversely, several natural and engineered nanobodies exhibit antigen-dependent stabilization, being destabilized and degraded when unbound, but stabilized upon engagement with their target (*35–39*). In a seminal study, Tang and colleagues used random mutagenesis and screening to identify destabilizing mutations in the GFP nanobody that confer antigen-dependent stabilization (*35*). These mutations are located in the conserved Framework Regions (FR) of the nanobody and can be transferred to other nanobodies to confer increased antigen-dependent stability (*35, 40*). Building on this strategy, we recently engineered a destabilized ALFA chromobody and demonstrated its utility for live imaging of an ALFA-tagged protein in drosophila (*9*). Nevertheless, the broader application of destabilized fluorescent binders for visualization of endogenous proteins has remained largely unexplored in multicellular systems.

Here, we develop a library of fluorescent protein binders for cell type-specific visualization of proteins fused to GFP, ALFA, V5, 127D01, gp41 peptide (MoonTag), GCN4 peptide (SunTag), HA or FLAG epitopes. The binders can be expressed from *UAS* transgenes for use with Gal4 driver lines in drosophila (*41*) or from tissue-specific promoters in zebrafish. In addition, we incorporated the previously described destabilized GFP nanobody (*35*) and engineered destabilized binder variants of the GFP, ALFA, V5 and GCN4 peptide (SunTag) binders that undergo antigen-dependent stabilization. We show that reducing the abundance of unbound binders using this strategy substantially improves the signal-to-noise ratio for endogenous protein visualization by decreasing background fluorescence while preserving target-associated labelling. We demonstrate that this strategy enables cell type-specific visualization of endogenous proteins *in vivo*, in drosophila and zebrafish embryos, as well as in defined neuronal populations of the adult drosophila brain. The library presented here will be of general use for cell-type labelling of endogenous proteins.

## Results

### Characterization of GFP chromobodies for cell type-specific protein visualization

GFP chromobodies have enabled the recolouring of GFP-tagged proteins with spectrally distinct fluorescent proteins in cell culture (*12*). To adapt this strategy for cell type-specific visualization of broadly expressed GFP-tagged proteins, we generated two chromobody constructs: GFPNb:mSc3, comprising the GFP nanobody (GFPNb) (*12*) fused to mScarlet3 (mSc3) (*42*), and dGFPNb:mSc3, in which the destabilized GFP nanobody (dGFPNb) (*35*) was fused to mSc3. Both constructs were placed under the control of 5x *UAS* sequences to enable expression in defined cell populations using different Gal4 driver lines with tissue-or cell type-specific expression patterns. We first tested target binding and antigen-dependent stabilization in drosophila wing imaginal disks. Using *hh*-Gal4, GFPNb:mSc3 or dGFPNb:mSc3 were expressed in the posterior compartment in the presence or absence of membrane-bound mCD8:GFP (*43*), which was expressed in the dorsal compartment via *ap*-LexA and *LexAop*-mCD8:GFP (*44*) (Figure 1A-E, Figure S1A). Both chromobodies colocalized with mCD8:GFP at the plasma membrane in the posterior dorsal compartment, confirming target binding (Figure 1C,D). In the absence of mCD8:GFP in the anterior ventral compartment, GFPNb:mSc3 produced a diffuse cytoplasmic and nuclear signal (Figure 1C), which was strongly reduced when expressing dGFPNb:mSc3 (Figure 1D,E). These observations indicate that the destabilizing mutations promote degradation of the unbound chromobody in drosophila while preserving antigen binding, consistent with previous findings in mammals (*35*).

**Figure 1.**
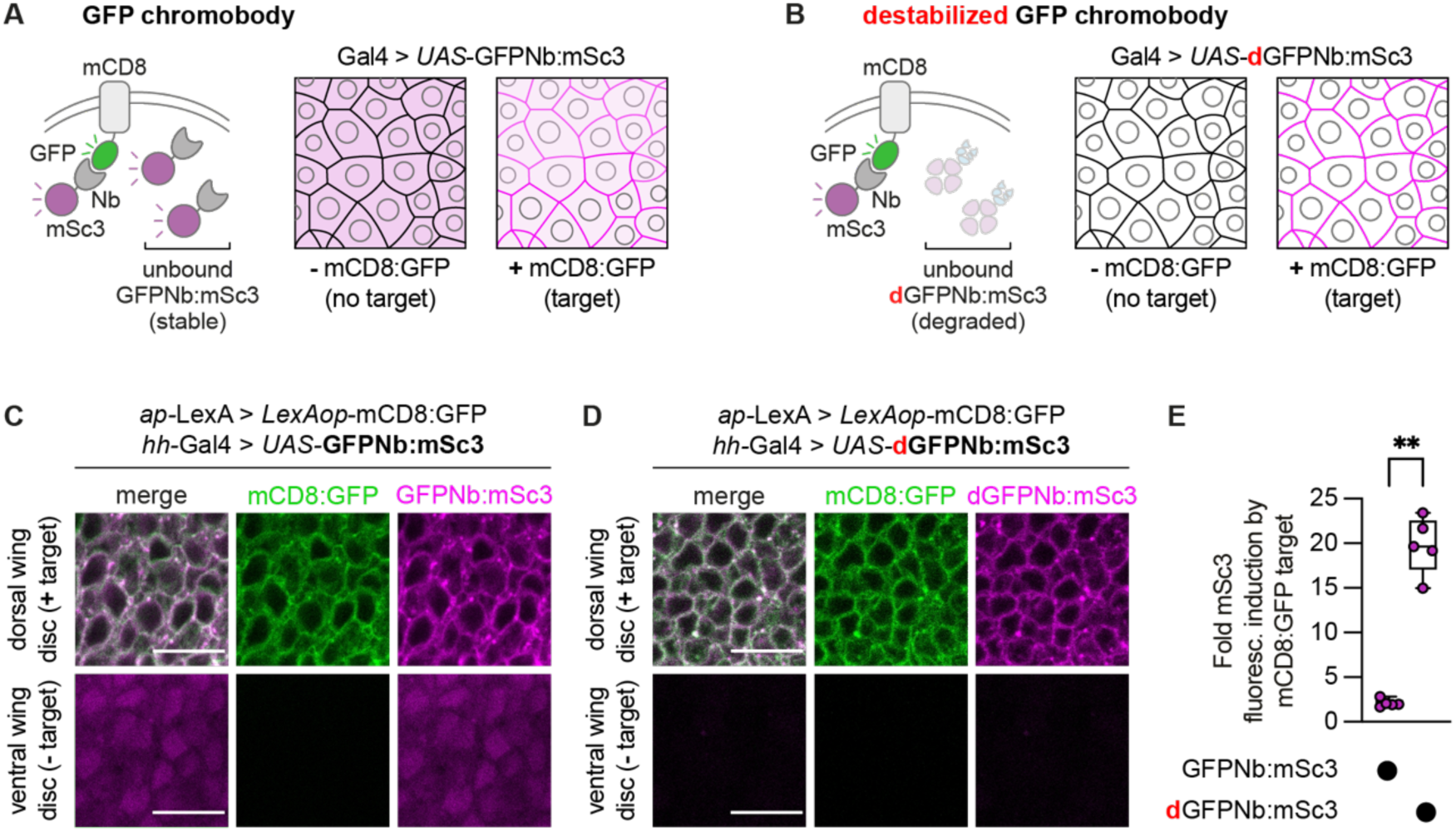
*In vivo* characterization of GFP chromobodies. **(A)** Schematic of GFP chromobody (GFPNb:mSc3)-mediated protein labelling. **Left:** GFPNb:mSc3 binds GFP fused to a target protein (here, mCD8) and labels it with mScarlet3 (mSc3). Unbound chromobody remains fluorescent and contributes to non-specific background signal. **Right:** Predicted distribution of Gal4-driven *UAS*-GFPNb:mSc3 in the absence or presence of mCD8:GFP. In the absence of the target, the chromobody is detected throughout the cell; in its presence, it accumulates at the plasma membrane while unbound chromobody remains diffuse. **(B)** Schematic of destabilized GFP chromobody (dGFPNb:mSc3)-mediated protein labelling. **Left:** dGFPNb:mSc3 binds GFP fused to a target protein (here, mCD8) and labels it with mSc3. Unbound chromobody is degraded, reducing background fluorescence. **Right:** Predicted distribution of Gal4-driven *UAS*-dGFPNb:mSc3 in the absence or presence of mCD8:GFP. In the absence of the target, little or no chromobody is detected; in its presence, binding to the target stabilizes the chromobody and results in its accumulation at the plasma membrane with minimal background signal. **(C and D)** Confocal microscopy images of wing imaginal disc cells expressing GFPNb:mSc3 (**C**) or dGFPNb:mSc3 (**D**) in the presence (top panels) or absence (bottom panels) of the mCD8:GFP target protein. (**E**) Fold mSc3 fluorescence induction of GFPNb:mSc3 and dGFPNb:mSc3 by the mCD8:GFP target. Fluorescence values were normalized to the mean intensity of the GFPNb:mSc3 or dGFPNb:mSc3 condition in the absence of the mCD8:GFP target protein. Genotype effect: *P* = 0.0079, Mann-Whitney test, *N* = 5. Data are shown as box-and-whisker plots (min to max) with all points displayed. *N*, number of technical replicates. Scale bars: 10 µm. For detailed genotype descriptions, see Table S4.

### Antigen-dependent stabilization improves the visualization of endogenous GFP-tagged proteins *in vivo*

Having established that the destabilized GFP chromobody undergoes antigen-dependent stabilization, we next asked whether this property improves the cell type-specific visualization of endogenous GFP-tagged proteins. We first examined the junctional protein E-cadherin (Ecad) in tracheal cells of drosophila embryos. GFPNb:mSc3 or dGFPNb:mSc3 were expressed using the tracheal-specific *btl*-Gal4 driver in embryos carrying an endogenously-tagged *Ecad^GFP^* allele (*45*) (Figure 2 A,B). For both chromobodies, the mSc3 signal colocalized with Ecad:GFP at apical junctions of tracheal cells, but not in neighbouring epidermal cells, demonstrating cell type-specific recolouring of Ecad:GFP. However, while GFPNb:mSc3 was also detected in the cytoplasm, where Ecad:GFP was largely absent, dGFPNb:mSc3 was restricted to Ecad:GFP-positive junctions (Figure 2A,B). Notably, GFPNb:mSc3 produced a stronger signal than dGFPNb:mSc3. Consequently, the imaging settings (including laser power) were adjusted individually for each chromobody to accurately visualize endogenous protein localization. These results show that antigen-dependent stabilization improves the cell type-specific visualization of endogenous proteins by reducing background fluorescence while preserving target binding.

**Figure 2.**
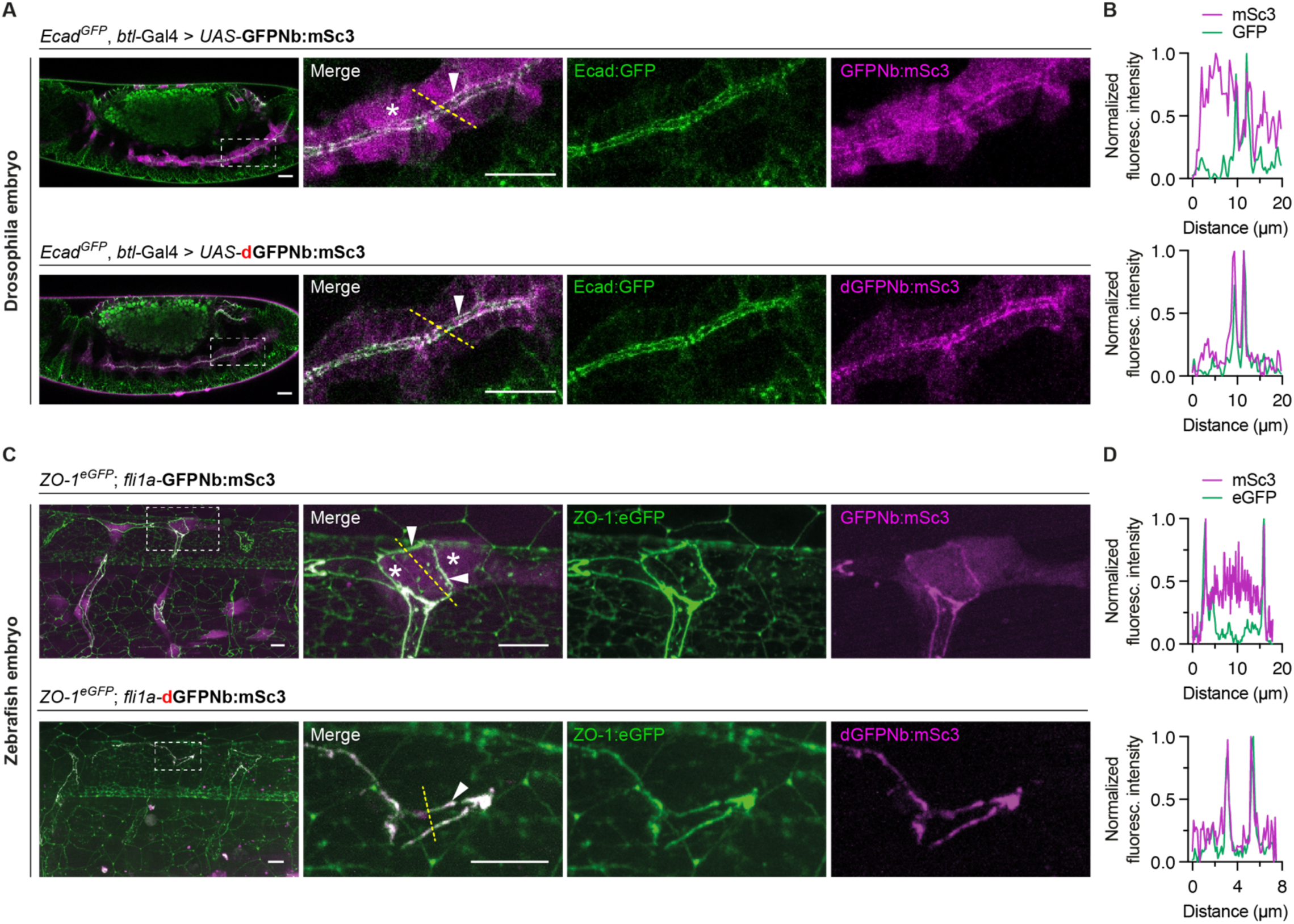
The destabilized GFP chromobody labels endogenous proteins with reduced background fluorescence. **(A)** Confocal live microscopy images of drosophila embryos (stage 14) expressing endogenous Ecad:GFP alongside GFPNb:mSc3 (upper panels) or dGFPNb:mSc3 (lower panels) in tracheal cells labelled using *btl*-Gal4. Insets display magnified views of tracheal cells, and arrowheads point to the region of the apical cell surface. The asterisk points to the GFPNb:mSc3 signal in the cytoplasm of tracheal cells. Scale bars: 20 µm. **(B)** Line-scan analyses of GFP and mSc3 fluorescence intensity along the dotted lines indicated in (A). **(C)** Confocal live microscopy images of zebrafish embryos showing the dorsal longitudinal anastomotic vessel and intersegmental blood vessels at 30 hpf. Endogenous ZO-1:eGFP is expressed alongside *fli1a*:GFPNb:mSc3 (upper panels) and *fli1a*:dGFPNb:mSc3 (lower panels). Insets display magnified views of endothelial cells. Arrowheads indicate regions in which ZO-1:eGFP colocalizes with GFPNb:mSc3 and dGFPNb:mSc3, respectively. The asterisk points to the GFPNb:mSc3 signal in the cytoplasm of endothelial cells. Scale bars: 10 µm. **(D)** Line-scan analyses of eGFP and mSc3 fluorescence intensity along the dotted lines indicated in (C). GFPNb:mSc3 produced a stronger fluorescent signal than dGFPNb:mSc3, requiring different imaging conditions for each chromobody to accurately visualize endogenous protein localization. For detailed genotype descriptions, see Table S4.

To determine whether these observations extend to proteins with different subcellular localizations, we next examined RacGAP50C, a cytokinesis-associated protein (*46–48*). In stage 11-15 embryos, endogenously tagged RacGAP50C:GFP (*49*) localized diffusely throughout the cytoplasm and nucleus and formed discrete puncta at cell-cell interfaces, as previously reported in the pupal notum (*49*) (Figure S2A,B). The puncta are consistent with RacGAP50C localization to midbody remnants connecting recently divided daughter cells, while the diffuse nuclear and cytoplasmic signal matches its previously described localization prior to cell division (*47*). Next, we expressed either GFPNb:mSc3 or dGFPNb:mSc3 using *hh*-Gal4 in drosophila embryos and assessed their ability to recolour RacGAP50C:GFP. Expression of either chromobody labelled RacGAP50C:GFP positive puncta (Figure S2A,B). However, only dGFPNb:mSc3 faithfully reproduced the diffuse nucleocytoplasmic GFP signal (Figure S2B), whereas strong cytoplasmic and nuclear background fluorescence from GFPNb:mSc3 interfered with the detection of the GFP signal (Figure S2A).

Having established that antigen-dependent stabilization can improve endogenous protein visualization in drosophila, we next asked whether this strategy could also be applied in vertebrates, where cell type-specific labelling of endogenous proteins via genetic methods is most challenging. We therefore examined the application of GFP chromobodies in zebrafish embryos using the endogenous junctional protein ZO-1:eGFP (*50*) during vascular development. We expressed GFPNb:mSc3 or dGFPNb:mSc3 under the control of the vascular endothelial promoter *fli1a* and examined the localization of the GFP chromobodies in embryos 30 hours post-fertilization (Figure 2C,D). We found that both chromobodies colocalized with ZO-1:eGFP at endothelial junctions of intersegmental blood vessels and dorsal longitudinal anastomotic vessels (Figure 2C,D). In embryos expressing GFPNb:mSc3, mSc3 fluorescence was also detected throughout the cytoplasm, whereas dGFPNb:mSc3 remained largely confined to endothelial junctions (Figure 2C,D), in line with the results obtained in drosophila. Due to the mosaic expression inherent to DNA injections, GFPNb:mSc3 was expressed at different levels across cells, resulting in considerable variation in fluorescence intensity both at junctions and in the cytoplasm. By contrast, dGFPNb:mSc3 exhibited more uniform fluorescence levels, consistent with antigen-dependent stabilization of the chromobody. As in drosophila, GFPNb:mSc3 produced a stronger signal than dGFPNb:mSc3, requiring a separate adjustment of the imaging conditions with each chromobody. Together, these experiments demonstrate that antigen-dependent stabilization can be applied across different model systems to improve endogenous protein visualization.

### GFP chromobodies enable cell type-specific protein labelling in the adult drosophila brain

Cell type-specific visualization of endogenous proteins is particularly important in the brain, where cells are densely packed and the fluorescence of the FP fusions from surrounding cells can obscure the subcellular localization patterns in the cells of interest. To determine whether antigen-dependent stabilization remains effective in the adult brain, we examined dorsal fan-shaped body (dFB) cells, whose axons project to the fan-shaped body of the adult drosophila central complex (Figure 3A). We first characterized the target-dependent stability of GFP chromobodies in dFB neurons by expressing GFPNb:mSc3 or dGFPNb:mSc3 together with mCD8:GFP using *R23E10*-Gal4 (*4, 44*). The mSc3 signal of both chromobodies colocalized with mCD8:GFP throughout axons, dendrites and cell bodies, confirming target binding (Figure 3B,C). In the absence of mCD8:GFP, GFPNb:mSc3 produced a diffuse background fluorescence that was most prominent in the cytoplasm and nuclei of cell bodies (Figure 3B). By contrast, background fluorescence was markedly reduced for dGFPNb:mSc3 in the absence of a target (Figure 3C,D, Figure S3A).

**Figure 3.**
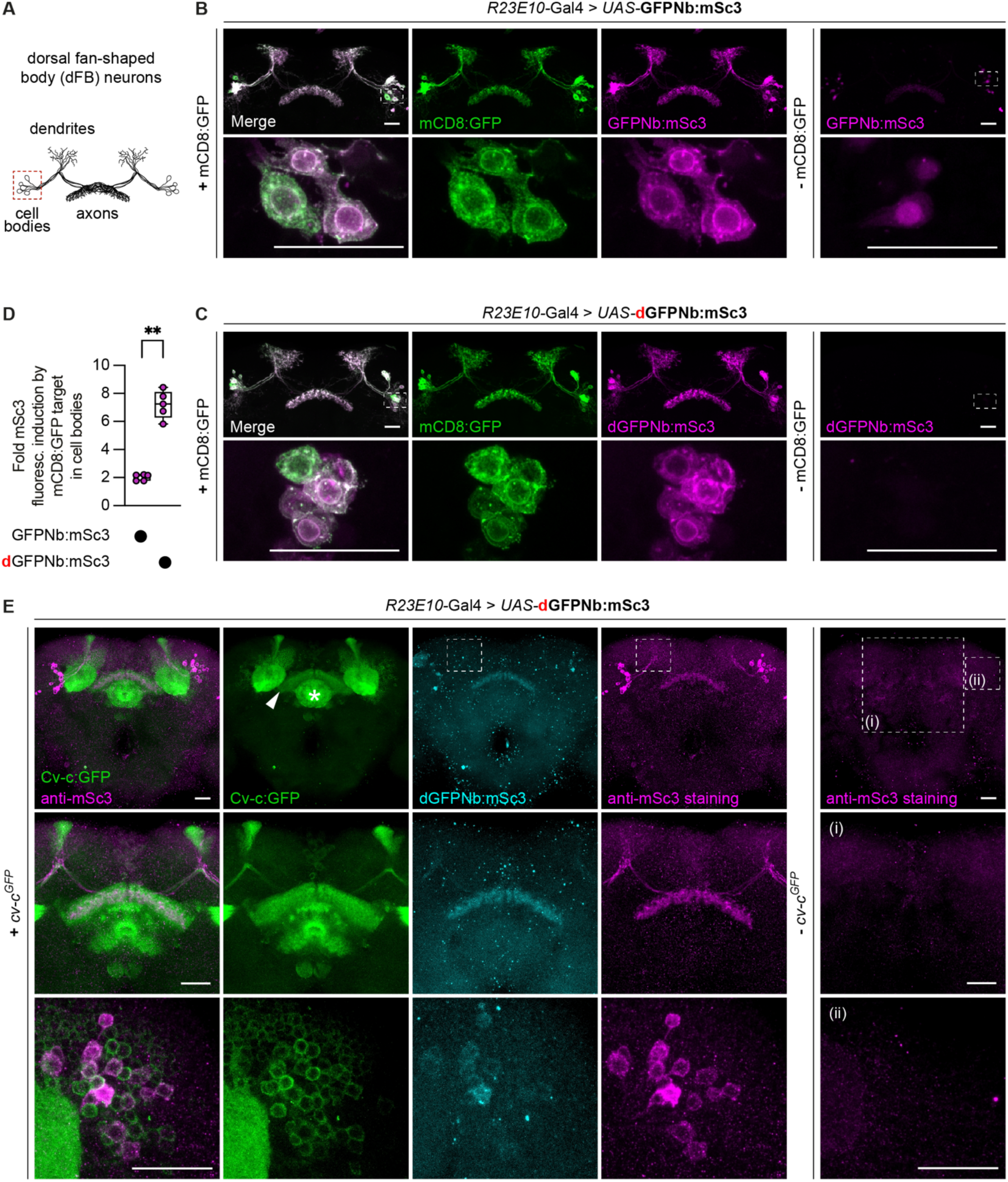
The destabilized GFP chromobody labels overexpressed and endogenous GFP fusion proteins in the adult drosophila brain. **(A)** Illustration of dendritic and axonal compartments of dorsal fan-shaped body (dFB) neurons labelled by *R23E10*-Gal4. **(B and C)** Confocal microscopy images of *R23E10*-Gal4 dFB neurons expressing GFPNb:mSc3 **(B)** or dGFPNb:mSc3 **(C)** in the presence (left) or absence (right) of mCD8:GFP. Dashed-line insets indicate selected cell bodies shown at a higher magnification in the lower panels. Scale bars: 25 µm. **(D)** Fold mSc3 fluorescence induction of GFPNb:mSc3 and dGFPNb:mSc3 by the mCD8:GFP target in cell bodies. Fluorescence values were normalized to the mean intensity of the GFPNb:mSc3 or dGFPNb:mSc3 condition in the absence of the mCD8:GFP target protein. Genotype effect: *P* = 0.0079, Mann-Whitney test, *N* = 5. Data are shown as box-and-whisker plots (min to max) with all points displayed. *N*, number of technical replicates. **(E)** Confocal microscopy images of brains expressing dGFPNb:mSc3 in dFB neurons in the presence (left) or absence (right) of Cv-c:GFP. The arrowhead points to the fan-shaped body and the asterisk to the ellipsoid body of the central complex. Dashed lines indicate the expected localization of the dendrites, which are not labelled. Magnified views show the fan-shaped body region (middle panels) and the cell body region (lower panels). Scale bars: 25 µm. For detailed genotype descriptions, see Table S4.

We next asked whether dGFPNb:mSc3 would also label proteins occupying other subcellular compartments. To this end, dGFPNb:mSc3 was expressed in dFB neurons using *R23E10*-Gal4 in the presence or absence of histones tagged with YFP (*ubi*-H2A:eYFP), which was shown to preserve the binding epitope of the GFP nanobody (*51, 52*). The chromobody-derived mSc3 signal colocalized exclusively with H2A:eYFP in neuronal nuclei with no detectable fluorescence in the cytoplasm (Figure S4A), or in the absence of H2A:eYFP (Figure S4B), confirming specific labelling of nuclear YFP-tagged proteins. Together, these results demonstrate that antigen-dependent stabilization enables specific labelling of GFP-tagged proteins in both membrane and nuclear compartments of adult neurons.

### Destabilized GFP chromobodies enable cell type-specific visualization of endogenous Cv-c in adult neurons

Having established that dGFPNb:mSc3 functions in adult neurons, we next asked whether it allows to visualize an endogenous GFP-tagged protein specifically in dFB neurons. We examined the Rho GTPase-activating protein (RhoGAP) Crossveinless-c (Cv-c), which plays a key role in the regulation of sleep homeostasis in these neurons (*53*). Transcriptomic data and dFB-restricted knockdown of *cv-c* support a role for Cv-c in sleep regulation, yet its protein expression and subcellular localization has not been reported (*53, 54*). In the adult brain, we found that endogenously tagged Cv-c:GFP (*49*) is expressed in axonal compartments of the mushroom body and parts of the central complex, including the dFB and ellipsoid body, as well as in neuronal cell bodies, where it localizes to the cell periphery (Figure 3E). Expression of dGFPNb:mSc3 in *R23E10*-Gal4 targeted dFB neurons carrying the endogenous Cv-c:GFP allele resulted in a faint but reproducible mSc3 signal in axons and cell bodies but not in dendrites, recapitulating the compartment-specific localization of endogenous Cv-c:GFP (Figure 3E). To increase detection sensitivity, we immunostained the chromobody using an anti-mCherry antibody, which enhanced the signal while preserving its localization (Figure 3E). Within neuronal cell bodies, the amplified chromobody signal colocalized with Cv-c:GFP at the cell periphery, faithfully reproducing its endogenous subcellular localization (Figure 3E). Thus, antibody amplification increased detection sensitivity while preserving the compartment-specific localization of endogenous Cv-c, including its exclusion from dendrites. Neither mSc3 fluorescence nor anti-mSc3 staining was detected in the absence of Cv-c:GFP (Figure 3E), confirming that chromobody localization depends on target binding. Together, these results establish destabilized GFP chromobodies as a strategy for visualizing endogenous neuronal proteins while preserving their subcellular localization.

### Engineering chromobodies directed against short epitope tags

To extend our approach beyond GFP, we next asked whether antigen-dependent stabilization could be transferred to nanobodies directed against a panel of commonly used short epitope tags. These tags permit the visualization of endogenous proteins with minimal modification of the genomic locus while remaining compatible with genetically encoded protein binders. We recently engineered a destabilized ALFA nanobody (dALFANb) (*9*) by introducing 5 of the 6 destabilizing framework mutations originally described for the GFP nanobody (*35*), while excluding the mutation located within the ALFA peptide-binding interface (*24*) (Figure 4A,B). The resulting dALFANb was fused to either Cherry or mGreenLantern to generate destabilized ALFA chromobodies (*9*). These ALFA chromobodies retained target specificity while exhibiting reduced background fluorescence in the absence of the target. Here, we applied the same engineering strategy to the V5 nanobody (V5Nb) (*25*), the 127D01 nanobody (127D01Nb) (*22*), and to the 2H10 gp41 nanobody (gp41Nb) (*26*). Destabilizing mutations were introduced into conserved framework residues while avoiding positions adjacent to the predicted peptide-binding interface, generating dV5Nb, d127D01Nb and dgp41Nb (Figure 4A,B, Figure S5J). Wild-type and destabilized nanobodies, including the previously described ALFANb and dALFANb, were fused to mSc3 to generate chromobodies. Thus, we generated a panel of chromobodies to systematically assess whether antigen-dependent stabilization could be transferred to distinct peptide-binding nanobody scaffolds.

**Figure 4.**
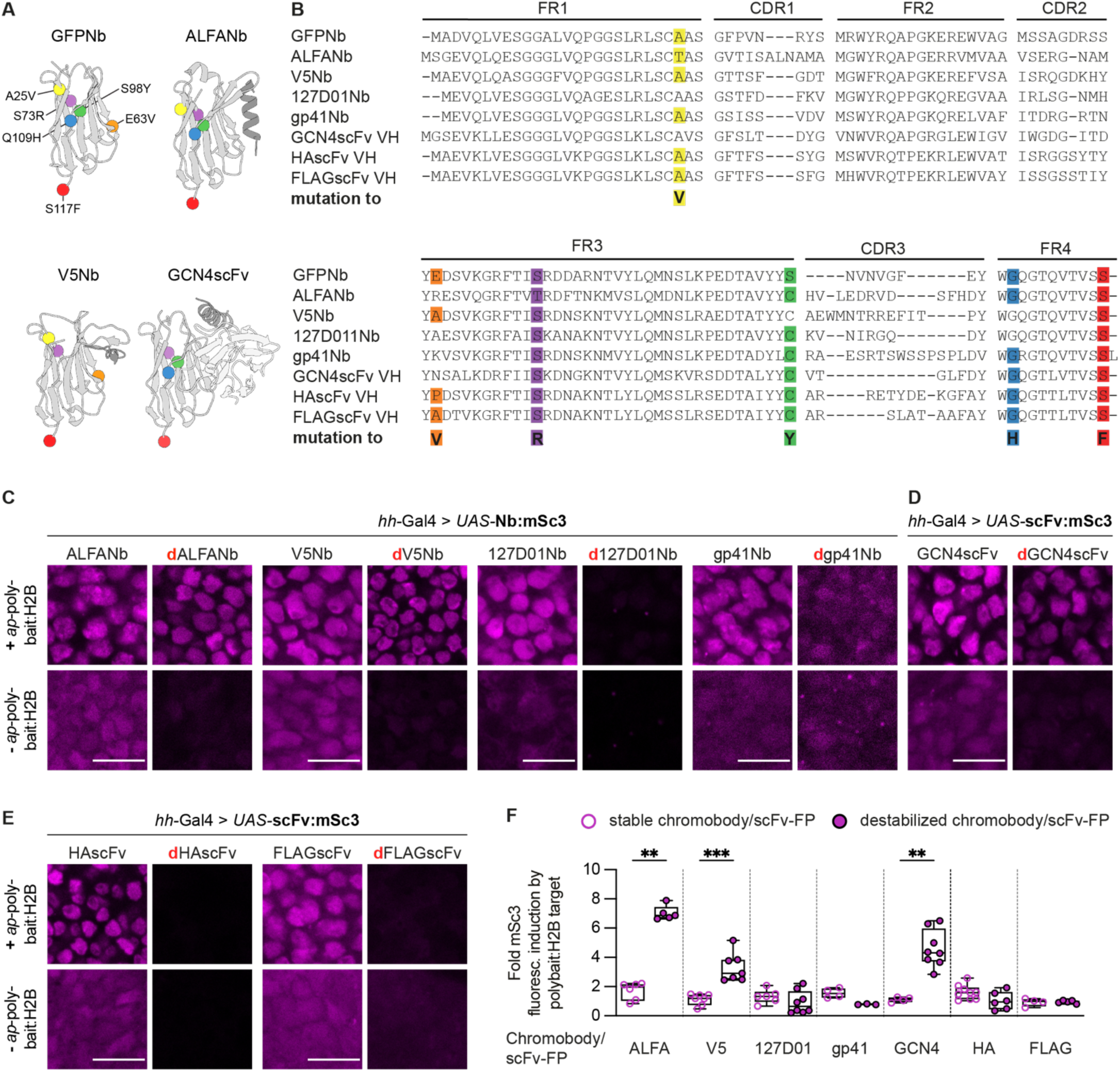
Engineering chromobodies and scFv-FP fusions directed against short epitope tags. **(A)** Structures of the GFP nanobody (PDB: 3OGO), ALFA nanobody with ALFA peptide (PDB: 6I2G), and V5 nanobody with V5 peptide (PDB:8SKJ), and structural model of the GCN4 scFv with GCN4 peptide (AlphaFold 3). Coloured spheres indicate the positions of the destabilizing substitutions. **(B)** Protein sequence alignment of nanobodies and scFv heavy chains (scFv VH). Destabilizing substitutions are color-coded as in (A). **(C-E)** Confocal microscopy images of dorsal wing imaginal disc cells expressing the chromobodies or scFv-FP fusions ALFANb:mSc3 or dALFANb:mSc3 (C), V5Nb:mSc3 or dV5Nb:mSc3 (C), 127D01Nb:mSc3 or d127D01Nb:mSc3 (C), gp41Nb:mSc3 or dgp41Nb:mSc3 (C), GCN4scFv:mSc3 or dGCN4scFv:mSc3 (D), HAscFv:mSc3 or dHAscFv:mSc3 (E), and FLAGscFv:mSc3 or dFLAGscFv:mSc3 (E) under the control of *hh*-Gal4 and in the presence (upper panels) or absence (lower panels) of polyBait:H2B (*ap*-polyBait:H2B). **(F)** Fold mSc3 fluorescence induction of chromobodies and scFv:mSc3 fusions by polyBait:H2B. Fluorescence values were normalized to the mean intensity of the respective stable or destabilized conditions in the absence of the polyBait:H2B target protein. Genotype effect: *P* = 0.0043 (ALFA), *P* = 0.0006 (V5), *P* = 0.0040 (GCN4), Mann-Whitney test, *N* = 3-9. Data are shown as box-and-whisker plots (min to max) with all points displayed. *N*, number of technical replicates. Scale bars: 10 µm. For detailed genotype descriptions, see Table S4.

To characterize target recognition and antigen-dependent stabilization, the chromobodies were expressed using *hh*-Gal4 in third instar wing imaginal discs, both in the presence or absence of polyBait:H2B (Figure S5A), a nuclear Histone H2B fusion protein carrying a peptide array containing ALFA, HA, V5, FLAG, 127D01, GCN4 peptide and gp41 peptide tags separated from each other by flexible linkers (Figure S5I). To approximate endogenous protein abundance, polyBait:H2B was expressed under the control of endogenous *apterous* (*ap*) regulatory sequences (see Materials and Methods). Wild-type chromobodies accumulated in nuclei expressing polyBait:H2B, indicating target binding, but also exhibited substantial cytoplasmic and nuclear fluorescence in the absence of the target (Figure 4C-E). Background fluorescence was highest for ALFANb:mSc3 and V5Nb:mSc3, where total fluorescence levels approached those observed in the presence of the target (Figure 4C, Figure S5E,F). In contrast, dALFANb:mSc3 and dV5Nb:mSc3 retained nuclear localization in the presence of polyBait:H2B while exhibiting markedly reduced fluorescence in its absence, resulting in a significant target-dependent increase in fluorescence (Figure 4F, Figure S5E,F). These findings indicate that antigen-dependent stabilization can be successfully transferred to the ALFA and V5 nanobodies. By contrast, d127D01Nb:mSc3 and dgp41Nb:mSc3 exhibited low fluorescence irrespective of target expression and strong background fluorescence in the cytoplasm (Figure 4C), indicating that the introduced framework mutations were interfering with antigen binding. Additionally, we observed the formation of small fluorescent foci for d127D01:mSc3 and dgp41Nb:mSc3 (Figure 4C), indicating altered behaviour of these destabilized fluorescent protein binders.

Together, these experiments establish a panel of chromobodies targeting ALFA, V5, 127D01 and gp41 peptide tags, and demonstrate the successful generation of destabilized ALFA and V5 chromobodies. To further expand the usefulness of the toolkit, we generated additional versions of dALFANb fused to mTagBFP2 (*55*) and Dendra2 (*56*), and dV5Nb fused to tandem dimer StayGold (tdSG) (*57*) (Table 1).

**Table 1.** Panel of validated fluorescent protein binders.

|  | **Fluorescent protein binder** |  |
| --- | --- | --- |
| **Target epitope** | **Wild type sequence binder** | **Destabilised binder** |
| GFP | GFPNb:mSc3 | dGFPNb:mSc3 |
| ALFA tag | ALFANb:mSc3 | dALFANb:mSc3<br>dALFANb:mGL<br>dALFANb:mtagBFP<br>dALFANb:dendra2 |
| V5 tag | V5Nb:mSc3 | dV5Nb:mSc3<br>dV5Nb:tdSG |
| 127D01 tag | 127D01Nb:mSc3 | d127D01Nb:mSc3 |
| gp41 peptide tag | gp41Nb:mSc3 | N/A |
| GCN4 peptide tag | GCN4scFv:mSc3 | dGCN4scFv:mSc3<br>dGCN4scFv:tdSG |
| HA tag | HAscFv:mSc3 | N/A |
| FLAG tag | FLAGscFv:mSc3 | N/A |

### Engineering scFv-FP fusions directed against short epitope tags

To further expand the repertoire of fluorescent protein binders recognizing short peptide tags, we next asked whether antigen-dependent stabilization could be transferred from nanobody to single-chain variable fragment (scFv) binders. We selected scFvs recognizing the GCN4 (SunTag) (*27, 58*), HA (*28*) and FLAG (*30*) epitopes, which have previously been used for protein labelling in model organisms (*31, 59*). We first generated GCN4scFv:mSc3, HAscFv:mSc3 and FLAGscFv:mSc3 binders (Figure 4A, Figure S5J). These fusion proteins were all robustly detected in nuclei expressing the polyBait:H2B target (Figure 4D,E). As observed for the nanobody-based chromobodies, however, all three constructs also exhibited cytoplasmic and nuclear fluorescence in the absence of the target, with FLAGscFv:mSc3 showing the highest background signal (Figure 4E, Figure S5D).

We next asked whether antigen-dependent stability could be transferred from nanobodies to the scFv scaffold. scFvs are composed of the immunoglobulin variable regions of heavy chain (VH) and light chain (VL), connected by a short peptide linker. Because the heavy chain of scFvs is structurally related to the nanobody scaffold, we hypothesised that nanobody-destabilizing mutations could be transferred into conserved framework residues of the VH domains of GCN4scFv, HAscFv and FLAGscFv, generating dGCN4scFv, dHAscFv and dFLAGscFv, respectively (Figure 4A,B). All three destabilized variants exhibited reduced fluorescence in the absence of polyBait:H2B, indicating successful destabilization. However, only dGCN4scFv:mSc3 retained nuclear accumulation in the presence of polyBait:H2B, demonstrating that it preserved antigen binding. In contrast, dHAscFv:mSc3 and dFLAGscFv:mSc3 failed to accumulate in target-expressing nuclei, indicating that the introduced mutations abolished binding to their respective epitopes (Figure 4E,F, Figure S5B-D). To expand the available fluorophore palette, we additionally fused dGCN4scFv to tandem dimer StayGold (tdSG) (Table 1). Together, these results demonstrate that the antigen-dependent stabilization can be successfully transferred to an scFv, although its effectiveness depends on the structural properties of the individual scFv.

### *In vivo* visualization of endogenous short epitope-tagged RacGAP50C

Having established that antigen-dependent stabilization can be transferred to nanobody- and scFv-based fluorescent protein binders recognizing short epitope tags, we next asked whether these binders could visualize an endogenously short epitope-tagged protein *in vivo*. To that end, we examined RacGAP50C, whose localization we had previously characterized using the endogenous GFP fusion protein (RacGAP50C:GFP, Figure S2). Using SEED-Harvest technology (*9*), we generated an endogenously triple-tagged RacGAP50C:ALFA:V5:GCN4 fusion protein (Figure 5A,B and Figure S6A,B) and tested whether its localization could be recapitulated using the corresponding destabilized fluorescent protein binders.

**Figure 5.**
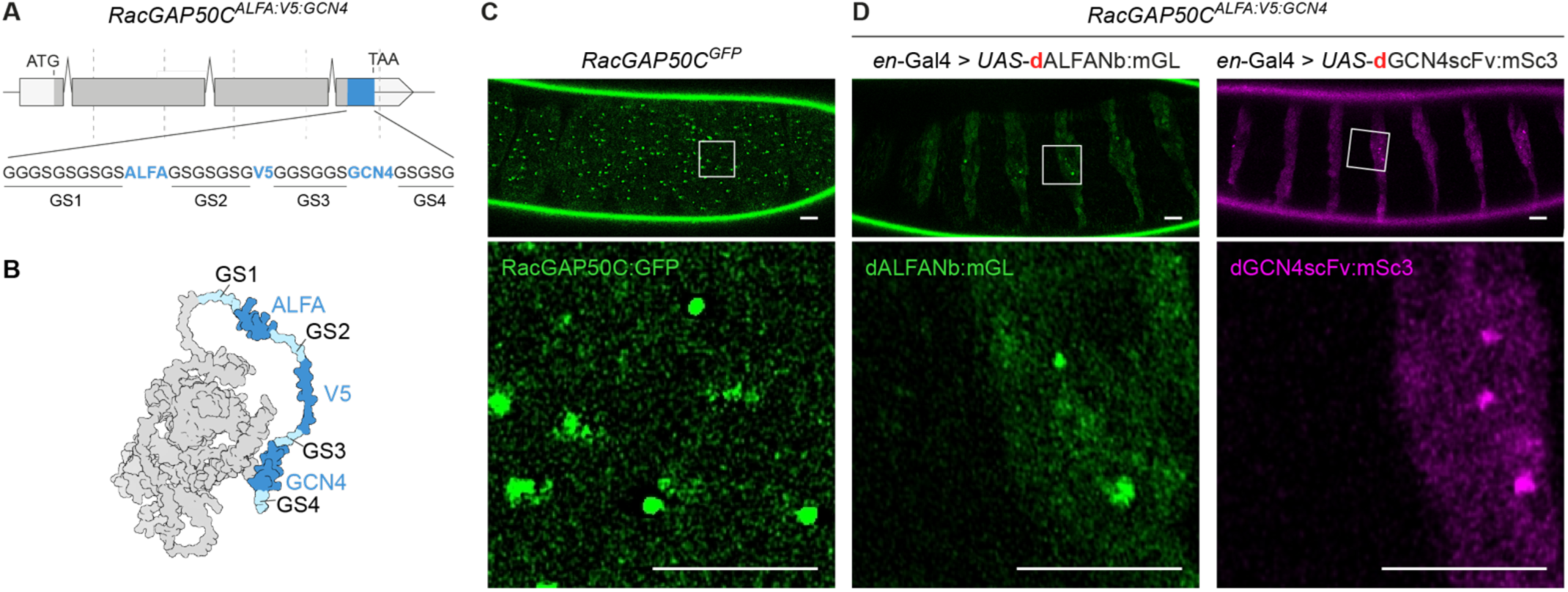
Chromobody labelling of short epitope-tagged RacGAP50C in drosophila embryos. **(A)** Schematic of the *RacGAP50C^ALFA:V5:GCN4^* allele carrying a C-terminal triple epitope tag composed of ALFA, V5 and GCN4 epitopes (blue) separated by flexible linkers (GS1-GS4). **(B)** Structural prediction of RacGAP50C:ALFA:V5:GCN4 with epitope tags highlighted in blue (AlphaFold 3 model). **(C and D)** Confocal live microscopy images of drosophila embryos (stages 14-15) expressing endogenous RacGAP50C:GFP **(C)**, or endogenous RacGAP50C:ALFA:V5:GCN4 alongside dALFA:mGL (left) or dGCN4:mSc3 (right) **(D)**. Insets show epidermal cells spanning the boundary between anterior and posterior segmental compartments. The left side of each inset corresponds to the anterior compartment, in which the chromobody is not expressed, whereas the right side corresponds to posterior-compartment epidermal cells labelled by *en*-Gal4. Scale bars: 10 µm. For detailed genotype descriptions, see Table S4.

Expression of dALFANb:mGL (Figure 5D), dGCN4scFv:mSc3 (Figure 5D) and dV5Nb:mSc3 (Figure S6C,D) in stage 14-15 embryonic epidermal cells produced diffuse nuclear and cytoplasmic fluorescence together with discrete puncta, closely matching the localization previously observed for RacGAP50C:GFP (Figure 5C). Together, these results establish tag-stabilized fluorescent protein binders as a versatile strategy for cell type-specific visualization of endogenous short epitope-tagged proteins *in vivo*.

## Discussion

Protein binders recognizing genetically encoded fluorescent proteins or short peptide epitopes have become versatile tools for visualizing and manipulating proteins in living cells and organisms (*10, 60*). Here, we developed a collection of tag-directed fluorescent protein binders for cell type-specific visualization of endogenous proteins. By incorporating the destabilized GFP nanobody (*35*) and engineering destabilized variants of ALFA-, V5- and GCN4-directed binders, we demonstrate that antigen-dependent stabilization can improve the signal-to-noise ratio of endogenous protein visualization, *in vivo* as well as in fixed tissues, by reducing background fluorescence while preserving target-specific labelling.

Our experiments show that the quality of protein binder-based labelling strongly depends on the balance between binder availability and target protein abundance. Highly abundant, ectopically expressed target proteins efficiently sequestered fluorescent binders, producing precise labelling with little interference from unbound molecules (Figure 1 C, D, Figure 3 B, D, Figure 4 C, D, E). By contrast, for endogenous proteins, where target abundance is lower, reducing binder stability markedly improved labelling by limiting the accumulation of unbound fluorescent binders. Thus, the relative abundance of target-bound and unbound binders is a key determinant of labelling specificity and of the accurate visualization of protein localization. Nanobodies used in cell biology are often soluble and stable in the cytoplasmic environment, while scFvs have been selected or engineered for these properties, enabling efficient target recognition but inevitably favouring the persistence of unbound binders (*27, 61, 62*). Several studies have shown that some binders display partial antigen-dependent stabilization even without engineering (*31, 39, 63*), probably because protein folding and antigen binding are thermodynamically coupled (*64*). In our study, however, this intrinsic antigen-dependent stability was insufficient to prevent substantial accumulation of unbound fluorescent binders.

Similar limitations have recently been reported for the visualization of endogenous ALFA-tagged Neurexin-1 in drosophila (*21*). To improve the signal-to-noise ratio by limiting the accumulation of unbound binders, we sought to make binder stability more responsive to antigen availability. Several approaches for engineering conditional stability have recently been described, including destabilizing framework mutations, split FP insertions and engineered N-terminal degron sequences (*35–39, 63, 65*). Our study demonstrates that destabilizing framework mutations substantially improve visualization of endogenous proteins *in vivo* by reducing background fluorescence while preserving target-dependent accumulation, allowing subcellular protein localization to be resolved in defined cell populations.

An important finding of this study is that antigen-dependent stabilization is not restricted to nanobodies but can be transferred to additional binder scaffolds. By introducing homologous framework mutations in the VH domain of the GCN4 scFv, we generated a destabilized variant that showed reduced fluorescence in the absence of antigen while retaining detectable binding to its cognate epitope. This result indicates that framework destabilization can be used to confer antigen-dependent stabilization beyond the single-domain nanobody format. The GCN4 scFv is widely used across fields for diverse applications, such as single-molecule imaging or transcriptional modulation (*27*). Further work will be needed to determine if antigen-dependent stability can improve the control of these applications.

Our analysis also shows that framework destabilization is not universally transferable between binders, consistent with previous reports (*35*). We found that destabilized variants of the 127D01 and gp41 nanobodies, as well as the HA and FLAG scFvs, abolished target protein binding, demonstrating that successful transfer of destabilizing mutations must be assessed empirically for each scaffold. The cytoplasmic puncta formed by the destabilized 127D01 and gp41 nanobodies further indicate that these variants display altered intracellular behaviour, probably aggregation due to excessive destabilization (Figure 4C). Importantly, the effect of destabilizing mutations may also depend on the experimental context. Antigen-dependent stability was previously not observed for a destabilized ALFA nanobody in HEK293T cells (*66*), indicating that target abundance, cellular context and experimental temperature may influence binder behaviour. Engineering antigen-responsive binders therefore requires tuning the balance between the unbound and antigen-bound states rather than simply increasing or decreasing overall stability. A complementary strategy is to enhance antigen responsiveness by accelerating the turnover of unbound binders through the N-end rule. Keller and colleagues used N-terminal destabilizing residues to reduce basal chromobody levels and increase the relative stabilization induced by antigen binding (*63*), while the more recent ANGEL system applied an N-terminal arginine degron to promote degradation of unbound NbALFA and its accumulation in the presence of the ALFA tag (*39*).

Our tag-stabilized protein binders enable precise determination of the expression and localization of endogenous proteins in defined cell types. Using Cv-c as an example – for which expression pattern and subcellular localization in the adult drosophila brain was previously unavailable – we show that it is expressed in sleep-regulatory dFB neurons and is selectively localized to presynaptic axonal compartments. This illustrates how destabilized protein binders can generate new biological insights in addition to serving as imaging tools. Such applications are particularly valuable in neuroscience, where determining the subcellular localization of signalling molecules, receptors or ion channels within defined neuronal populations can provide important clues about neuronal circuit organization and computation.

The collection of orthogonal tag-binder systems described in this study will substantially expand the experimental flexibility of endogenous protein visualization and facilitate multiplexed visualization strategies. We demonstrate that destabilized GFP, ALFA, V5 and GCN4 binders faithfully label endogenous proteins in drosophila embryos and larvae, the adult brain and zebrafish embryos, illustrating that the strategy is applicable across tissues, developmental stages and animal models. In particular, the ability to visualize an endogenously triple-tagged RacGAP50C protein using three independent binders highlights the robustness of short peptide-tag strategies and provides a foundation for multiplexed endogenous protein imaging using orthogonal tag-binder pairs. The growing repertoire of genetically encoded peptide tags should further facilitate flexible experimental designs requiring simultaneous visualization of multiple proteins.

Our results revealed that antigen-dependent stabilization substantially improved signal specificity but also decreased target-associated fluorescence when compared with stable binders. This is likely because unbound binders are continuously degraded, reducing the total pool available for target binding. When the target is scarce, antigen-dependent stabilization may not fully compensate for binder loss. Consequently, visualization of low-abundance endogenous proteins required increased laser power or antibody amplification, as illustrated by endogenous Cv-c in adult neurons. Signal intensity could potentially be enhanced by increasing the number of epitope copies per target molecule using peptide-tag arrays (*33, 67*) or multivalent “spaghetti monster” scaffolds (*68, 69*), thereby recruiting multiple fluorescent binders while maintaining low background fluorescence.

Our findings also suggest that antigen-dependent destabilization could be useful beyond visualization, particularly when intracellular binders are coupled to functional effector domains. In such applications, constitutively stable binders carrying enzymatic or other active domains could produce off-target activities. Restricting effector abundance to antigen-positive cells could reduce off-target activity while improving the spatial precision of protein manipulation (*70*). The increasing availability of orthogonal peptide tag-binder pairs also raises the possibility of simultaneously visualizing and manipulating the same endogenous protein using distinct binders directed against adjacent epitopes. Such strategies have recently been proposed conceptually (*9*) but remain largely unexplored experimentally. For *in vivo* protein studies, tag spacing and compatibility of distinct functionalized protein binders will require systematic validation, but insertion of several short peptide epitopes certainly provides greater flexibility in the concomitant manipulation and visualization of proteins in their complex cellular environment. The toolkit presented here provides a foundation for developing multifunctional intracellular binders that combine protein visualization with conditional manipulation in living organisms.

Together, our findings establish tag-stabilized fluorescent protein binders as a versatile strategy for improving cell type-specific visualization of endogenous proteins *in vivo*. By reducing background fluorescence while remaining compatible with diverse nanobody and scFv scaffolds, antigen-dependent stabilization substantially expands the toolkit available for endogenous protein imaging. Beyond visualization, the transferability of this engineering strategy provides a framework for the future development of multifunctional intracellular binders capable of both monitoring and manipulating proteins in living organisms.

## Materials and Methods

### Drosophila strains and culture

Fly stocks were grown on standard polenta-agar-yeast fly food. Experiments were conducted at 25°C, 60% humidity. Table 1 lists the validated drosophila strains of fluorescent protein binders generated in this study. All drosophila strains used in this study are listed in the Key resource table (Table 2). Detailed descriptions of the generation of drosophila strains in this study are given below.

**Table 2.**
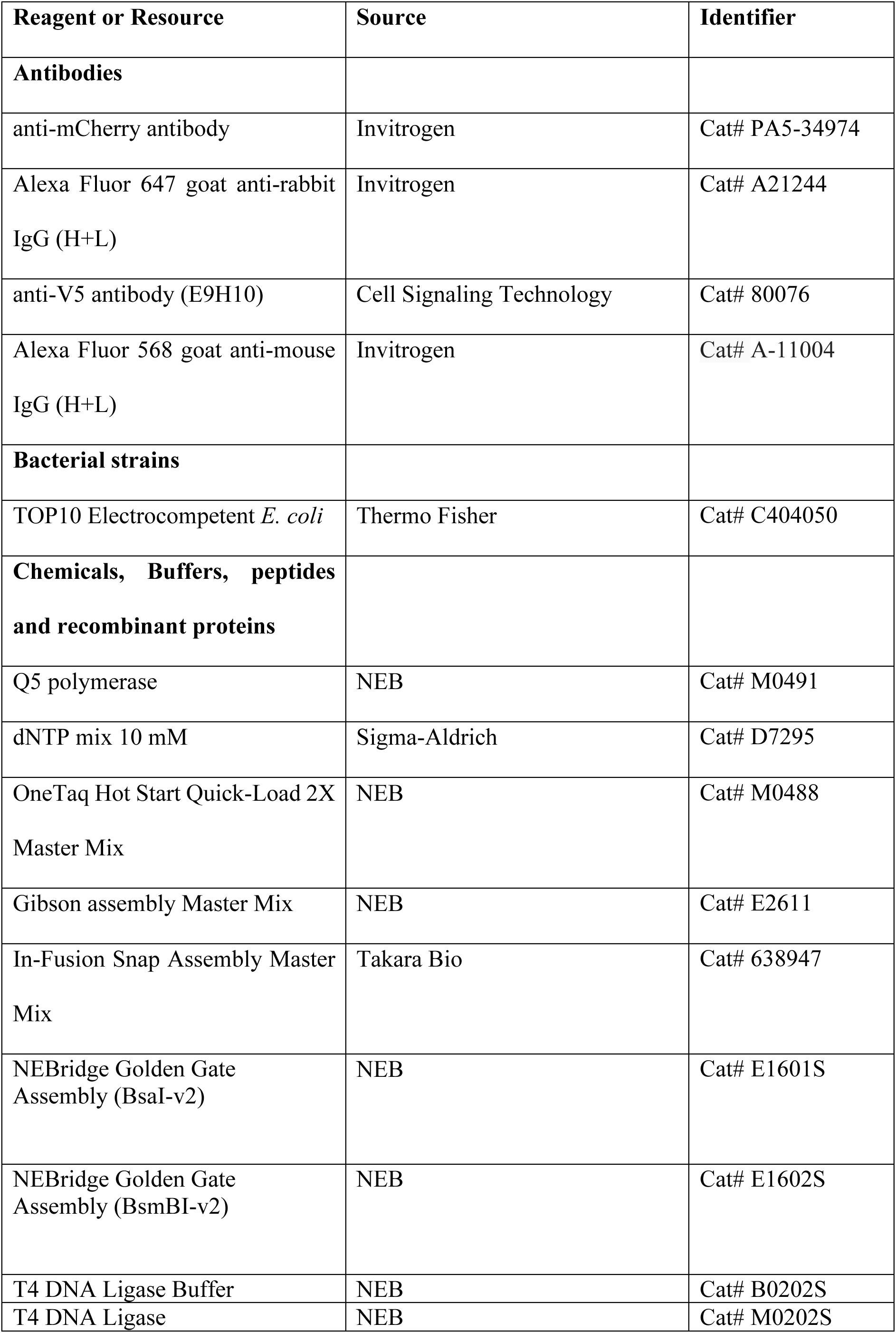
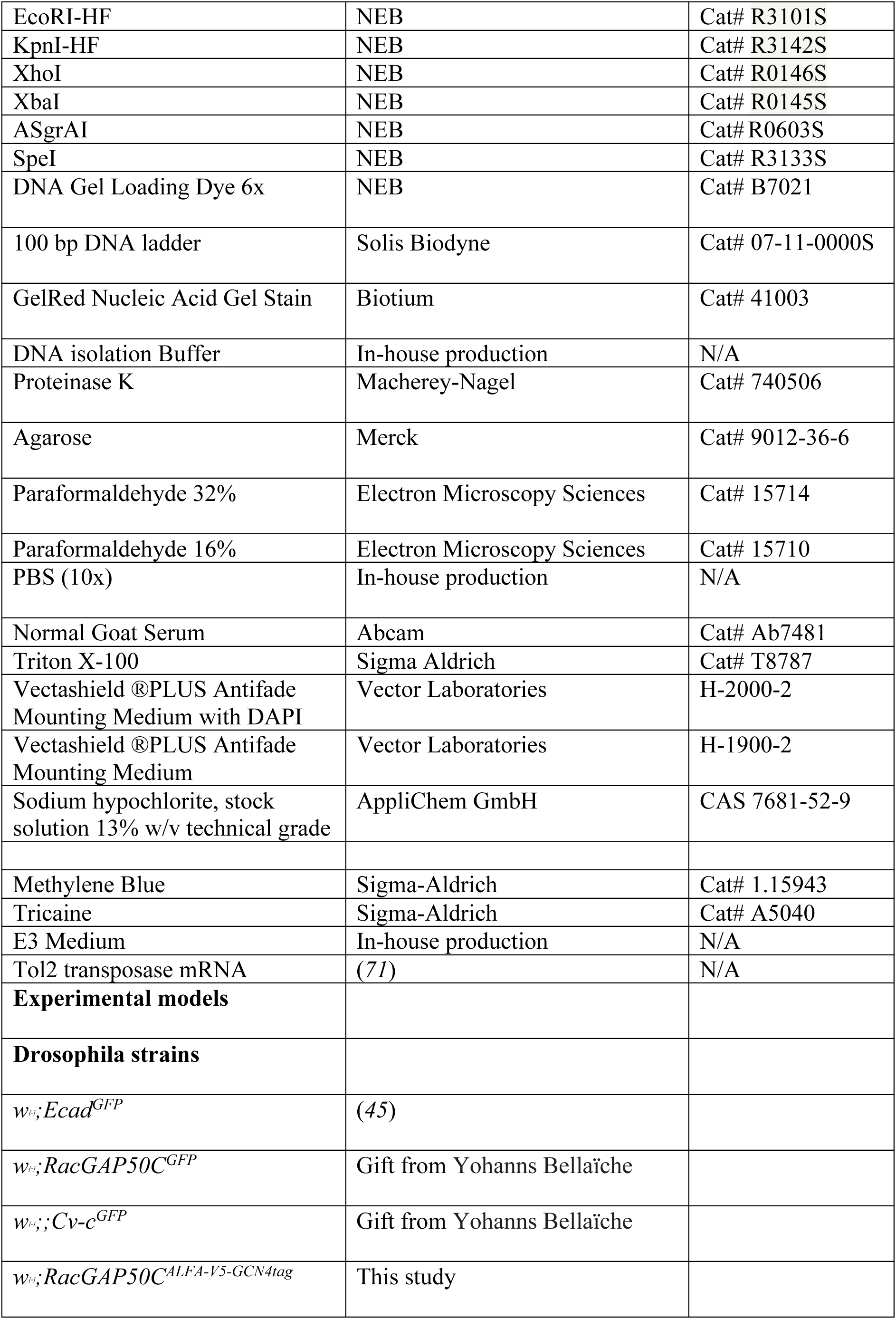
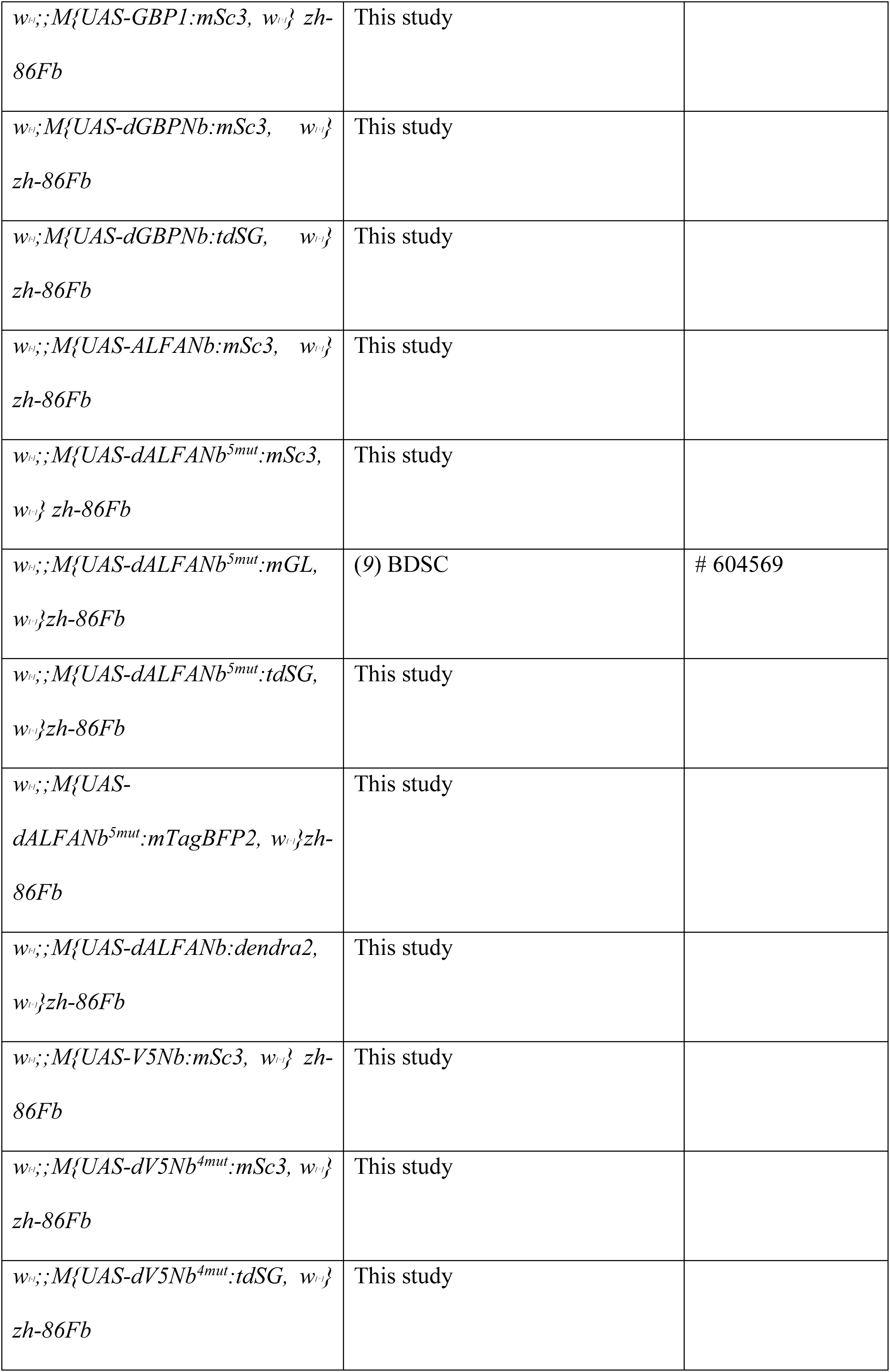
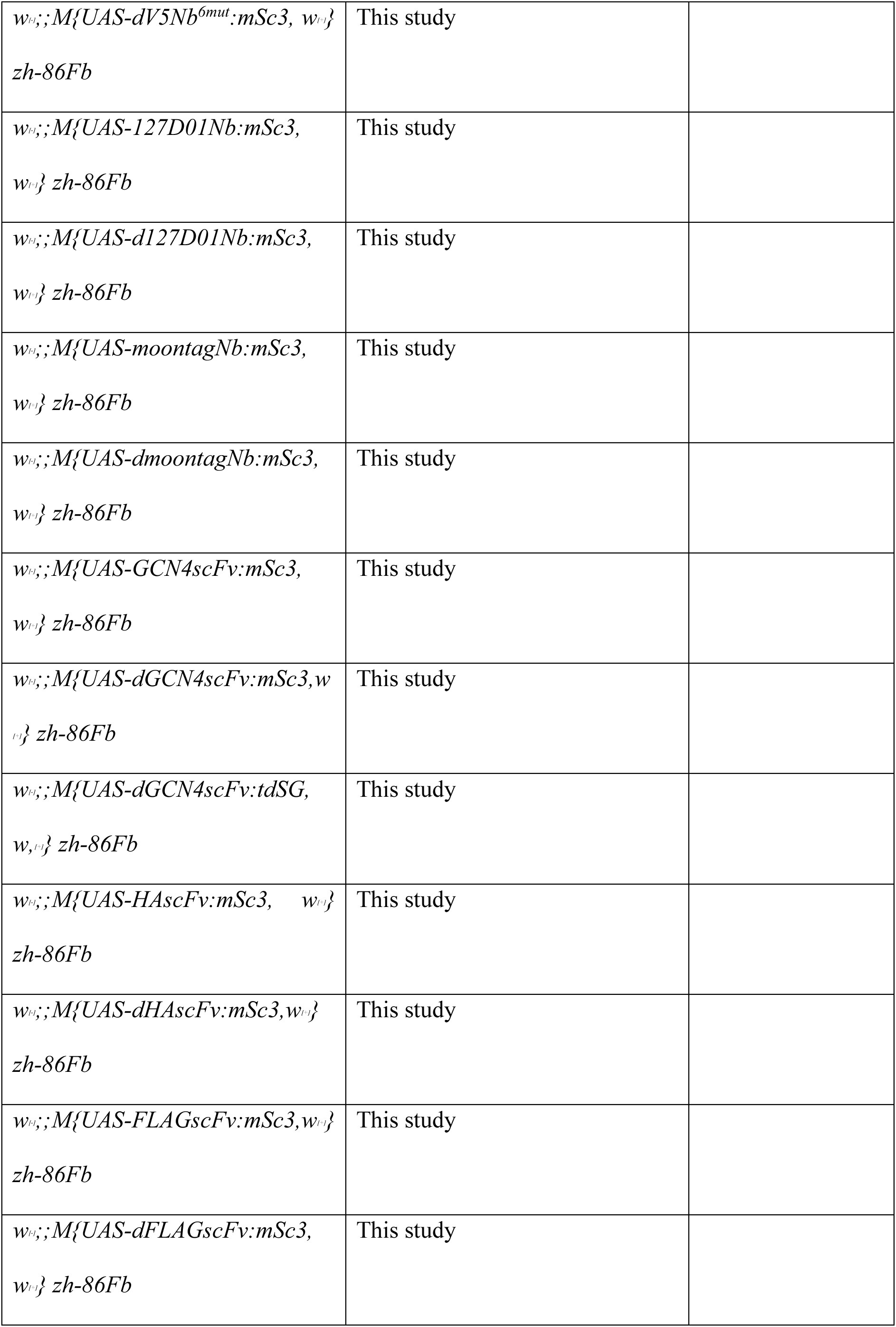
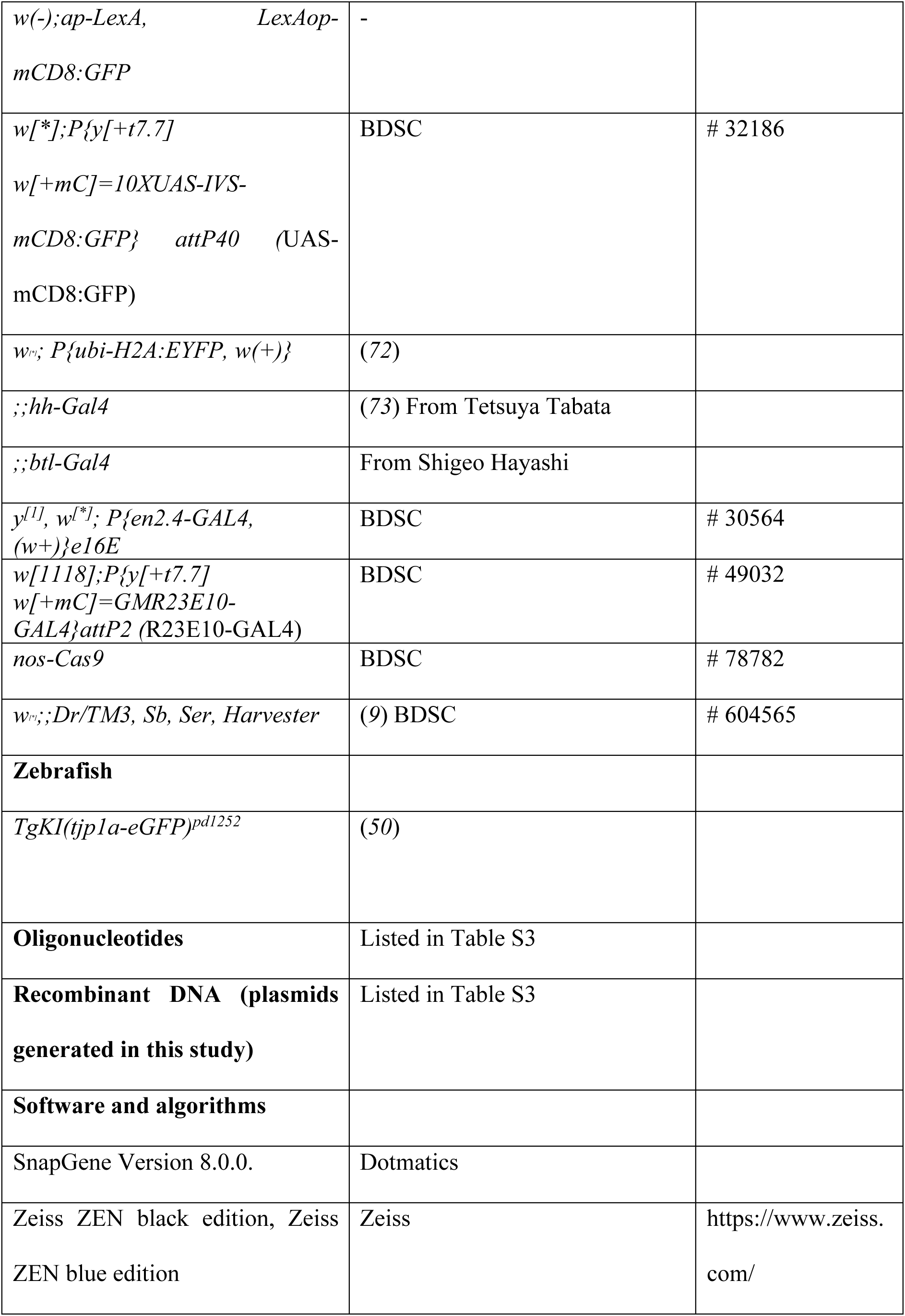
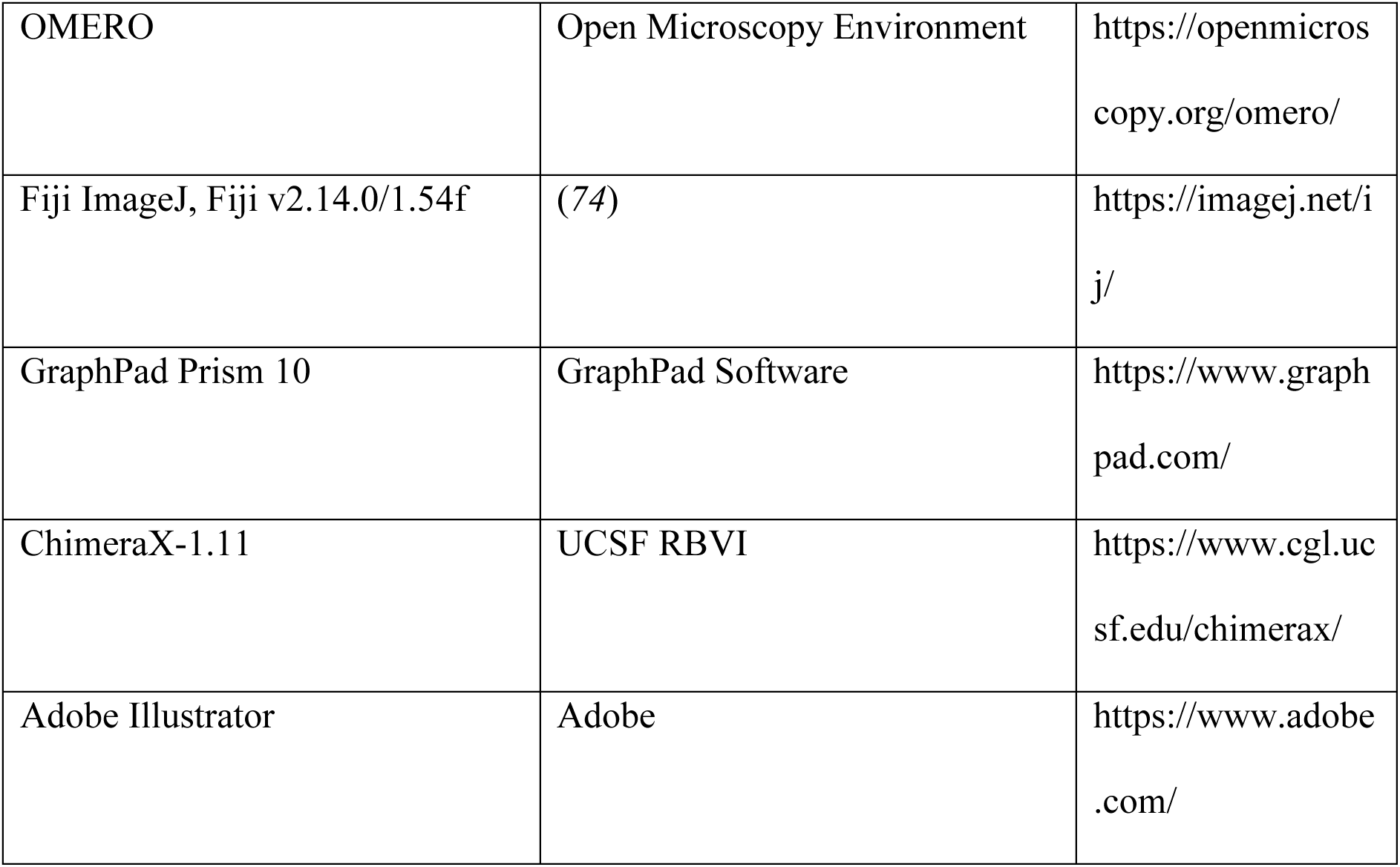
Key resource table.

### Molecular cloning

#### Generation of *pUAS_attB* fluorescent protein binders

DNA fragments for GFPNb (GBP1), dGFPNb (dGBP1), ALFANb, dALFANb^5mut^ (*9*), V5Nb, dVNb, MoonTagNb, dMoonTagNb, 127D01Nb, d127D01Nb, GCN4 scFv, dGCN4 scFv, HA scFv, dHA scFv, FLAG scFv, dFLAG scFv, mScarlet3, tdStayGold, mGreenLantern, mTagBFP2, Dendra2 (listed in Table S1) were purchased from Twist Bioscience or Genewiz as Gene Fragments. Nanobodies and scFvs were C-terminally fused with FPs spaced by a Serine-Glycine-Linker GGGSGGGS. Constructs – preceded by the Kozak sequence (ACCACC) – were assembled using Gibson Assembly in the pUASattB vector (*75*) linearized with EcoRI and KpnI. A list of the mutations in nanobodies and scFvs is provided in Table S2.

#### Generation of *polyBait:H2B*

Sequences containing part of ap regulatory sequences, the mhc intron, the polytag sequence, the coding sequence of H2B and the P74 sequence, were cloned into DB92-apReentry (*76*), using the AgeI and AscI restriction enzymes at the 5’ and 3’ ends, respectively. GGGS linkers were added between each tag, (GGGS)x2 was incorporated between the polytag and the H2B sequence.

#### Generation of fli1a:GFPNb:mSc3 and fli1a:dGFPNb:mSc3

GFPNb:mSc3 and dGFPNb:mSc3 were PCR amplified and ligated using In-Fusion cloning in fli1a:PH-AKT-EGFP (courtesy of Christian Helker) (*77*) linearized with ASgrAI and SpeI.

### Generation of transgenic drosophila lines

pUASattB plasmids containing fluorescent protein binders were injected with 300ng/ml DNA in PBS into embryos with attP ZH-86Fb landing site and nos-phiC31 following the embryo injection protocol described in (*9*). The ap-polyBait:H2B was generated by targeted insertion into the *ap ^c1.4b^,* as previously described (*76*).

### SEED/harvest knock-in RacGAP50C:ALFA-V5-GCN4

RacGAP50C:ALFA-V5-GCN4 C-terminal knock-in was generated following the protocol for the SEED/Harvest technology described in (*9*). RacGAP50C gRNA (5’-AATAACGAGACTTTCCTTGTTGG-3’) (Table S3) was designed with flyCRISPR and cloned into pCFD5 (Addgene 73914) by BbsI digestion and Gibson Assembly as described in (*78*). To generate the pSEED donor plasmid, RacGAP50C 180 bp homology arms flanked by gRNA target sites and spaced by the BsmBI cassette were synthesized by Twist Bioscience and assembled with the pSEED:ALFA-V5-GCN4 (with 3xP3dsRedx) using NEBridge^®^ Golden Gate Assembly BsmBI-v2. Donor vectors and gRNA plasmids were injected at 100 ng/µl in nos-Cas9 (3^rd^ Chromosome) flies. Injected flies were crossed with *y^[1]^, w^[-]^* and F1 progeny was screened for SEED integration using 3xP3dsRedx. The SEED cassette was harvested (3xP3dsRedx removed) using *w^[*]^;;Dr/TM3, Sb, Ser, Harvester.* Knock-ins after harvesting were confirmed by PCR and Sanger sequencing with primers: RacGAP50C_exon3_fwd: GATGAAGGCACTTCTTGAGCTGC; RacGAP50C_exon4_rev: GTTGGGTACAAATACAGCTCG; ALFAfwd: CGTTTGGAAGAGGAACTCAGAC; GCN4rev: GCCACTTCGTTCTCAAGATG.

### Evaluating destabilization and target binding of chromobodies and scFv-FP fusions in wing imaginal discs

Destabilization and target binding ability of chromobodies and scFv-FP fusions were evaluated in third-instar larvae wing disc cells. For GFP chromobodies, *ap-LexA, LexAop-CD8:GFP/CyO;hhGal4/TM6B* animals were crossed with *UAS-GFPNb::mSc3* or *UAS-dGFPNb:mSc3* animals. For chromobodies and scFv-FP fusions directed against short epitope tags a*p-polyBait:H2B/CyO; hh-Gal4/TM6B* animals or *if/Cyo;hh-Gal4/TM6B* (without target condition) animals were crossed to animals containing the UAS-chromobody or UAS-scFv-FP fusion. To isolate wing discs, larvae were dissected in ice-cold PBS and fixed in a paraformaldehyde solution (4% PFA in PBS) for 25 min at RT. Samples were rinsed 3 times for 5 min in PBS, and transferred to VECTASHIELD® PLUS Antifade Mounting Medium with DAPI. Wing discs were separated from the rest of the larvae, placed on a glass slide, and samples were covered with a coverslip and sealed using nail polish. Images of wing discs were acquired using Point Scanning Confocal Zeiss LSM880 (Carl Zeiss) with 40x/1.2 NA objective and silicon oil immersion, except for samples with ALFA chromobodies, which were acquired with Point Scanning Confocal Zeiss LSM700 (Carl Zeiss) with a 40×/1.3NA objective with oil immersion. For quantifications, mean fluorescent signal intensities were extracted from ROI squares of 32.95×32.95µm (Figure 1C,D and S1) and 41.53×41.53µm (Figure 4C-E and S5) of single planes of wing imaginal discs using Fiji ImageJ. For fold-change quantifications, values were normalized to the mean signal intensity of the wild-type or destabilized chromobodies or scFv-FP fusions in the absence of the target. For intensity quantifications, values were only normalized to the mean signal intensity of the wild-type chromobodies or scFv-FP fusions in the absence of the target. Values were plotted using GraphPad Prism Software. Image panels were prepared using Open Microscopy Environment (OMERO).

### Confocal live imaging of embryonic tracheal cells and embryonic epidermis

Embryos were dechorionated in 6% bleach and washed in deionized water. Embryos at the desired stage were manually selected on grape juice agar plates under a stereomicroscope, mounted on Glass Bottom MatTek Dishes, P35G-1.5-14-C, 35 mm petri dish, 14 mm microwell, and covered with 1xPBS. Tracheal and epidermal cells were imaged using the Point Scanning Confocal Zeiss LSM880 (Carl Zeiss) with a 40×/1.2NA objective (water) in confocal mode (Figure 2A) or AiryScan detector (Figure 5). Image panels were prepared using OMERO and fluorescent signal intensities were extracted using Fiji software and displayed using GraphPad Prism. For high resolution imaging of embryo features in Figure S2, the z-offset was corrected based on PSF measurements with fluorescent beads (Invitrogen).

### Immunostaining of drosophila embryos and image acquisition

Embryos were fixated according to standard protocol described in (*70*). Embryos were blocked with PBS with 0.5% Triton X-100, 2% NGS for 1 hour at RT, subsequently incubated with primary anti-V5 antibody 1:1000 in blocking solution overnight at 4°C, rinsed 3x for 20 min at RT in PBS with 0.5% Triton X-100 and incubated secondary Alexa Fluor™ 568 goat anti-mouse IgG H+L, Invitrogen, 1:1000 in blocking solution for 2 hours at RT. Embryos were rinsed 3x for 20 min at RT in PBS with 0.5% Triton X-100 and mounted in VECTASHIELD® PLUS Antifade Mounting Medium. Images were acquired with Point Scanning Confocal Zeiss LSM880 (Carl Zeiss) with 40x/1.2 NA objective and silicon oil immersion. Image panels were prepared using OMERO.

### Immunostaining of adult brains and image acquisition

To isolate brains, adult females (6-9 days after eclosion) were dissected in ice cold PBS. Samples were fixed for 20 min at RT in a paraformaldehyde solution (4% PFA in PBS with 0.5% Triton X-100) and rinsed three times for 20 min at RT in PBS with 0.5% Triton X-100. Samples were either mounted directly, or stained as follows: incubation for one day at 4°C in blocking solution (5% NGS in PBS with 0.5% Triton X-100), incubation for three days at 4°C in primary antibody solution (anti-mCherry rabbit polyclonal antibody, Invitrogen, 1:1000 in blocking solution), rinsing three times for 20 min at RT in PBS with 0.5% Triton X-100, incubation for two days at 4°C in secondary antibody solution (Alexa Fluor™ 647 goat anti-rabbit IgG H+L, Invitrogen, 1:1000 in blocking solution), rinsing three times for 20 min at RT in PBS with 0.5% Triton X-100. For mounting, the brain samples were transferred to 80% glycerol and placed on glass slides with their posterior side facing upward. Glycerol was removed, and VECTASHIELD PLUS Antifade Mounting Medium was added. Samples were covered with a coverslip and sealed using nail polish. Images were acquired with Point Scanning Confocal Zeiss LSM700 with a 20×/0.8NA objective (air) for images of full brains, and with a 40×/1.3NA objective with oil immersion for images of the regions of axons and cell bodies, and Point Scanning Confocal Zeiss LSM880 (Carl Zeiss) with 25x/0.8NA (oil) or 60×/1.2NA objective (silicone oil) for images in Figure S4. For quantification, Z-stacks of 20.4 µm containing only the dFB neuronal cell bodies from full brain acquisitions were flattened by *sum slices intensity projections*, and mean fluorescent signal intensities were extracted from ROI squares of 62.5×62.5 µm using Fiji ImageJ. Values were normalized to the mean signal intensity of the GFPNb:Sc3 or dGFPNb:Sc3 in the absence of the mCD8:GFP target and plotted using GraphPad Prism Software. Image panels were prepared using Open Microscopy Environment (OMERO).

### Zebrafish maintenance

Zebrafish (*Danio rerio*) were maintained according to FELASA guidelines (*79*). All experiments were performed following institutional and ethical welfare guidelines and animal protocols in accordance with federal guidelines approved by the Kantonales Veterinäramt of Kanton Basel-Stadt (1027H, 1014HE2, 1014 G). Breeding and embryo collection were done according to standard protocols (*80*). Pain, distress, and discomfort were minimized as much as possible.

### Expression of GFP chromobodies in zebrafish embryos

One-cell-stage embryos *TgKI(tjp1a-eGFP)^pd1252^* were injected with *fli1a*:GFPNb:mSc3 or *fli1a*:dGFPNb:mSc3. Plasmids were co-injected with Tol2 transposase mRNA at 35 ng/µl. Following injection, embryos were maintained at 28.5 °C in E3 medium containing 5 mg/l methylene blue to minimize fungal contamination. 30 hpf embryos were manually dechorionated and anesthetized in 1× E3 medium containing tricaine (1×). For imaging, embryos were embedded in 0.55% low-melting-point agarose, which was supplemented with tricaine, and mounted onto Glass Bottom MatTek Dishes. Embryos were positioned in a lateral orientation, with the anterior side facing left. After the agarose had solidified, samples were covered with E3 medium containing tricaine. Images were acquired with Olympus SpinSR (CSU-W1) using a 60X/1.3NA silicon oil objective, with a z-stack step size of 0.6 μm. 565 laser power was set to the mSc3 intensity levels of the GFPNb:mSc3 construct and dGFPNb:mSc3. Z-stacks were flattened by *maximum slice intensity projections* and image panels were created using the Open Microscopy Environment (OMERO).

### AlphaFold 3 modelling

gp41Nb-gp41 peptide, 127D01Nb-peptide, GCN4scFv-peptide, HAscFv-peptide and FLAGscFv-peptide structures were predicted with AlphaFold Server web interface (alphafoldserver.com). Binders and peptides were submitted as separate protein chains and default server settings. RacGAP50C:ALFA:V5:GCN4 engineered protein sequence was submitted with default server settings. Models were visualized with USCF Chimerax (v1.11).

## Acknowledgements and Funding

We thank Cindy Reinger and Matthias Zeug for fruitful discussions that helped move this project forward. We also thank Roman Jakob, Nikolaus Dietz, Timm Maier, Daan Overwijn and Maria Hondele for helpful advice. We thank the imaging core facility of the Biozentrum for their support on this project and Zofia Borzyszkowska for illustrations. We thank the Bloomington Stock Center for providing fly stocks. The work in the laboratory of M.A. was supported by grants from the Swiss National Science Foundation (310030_192659/1) and by funds from the Kanton Basel-Stadt and Basel-Land. S.T.S. was additionally supported by a ‘‘Biozentrum Fellowship” from the International PhD Program in Molecular Life Sciences of the Biozentrum, University of Basel. The work in the laboratory of A.K. was supported by the Swiss State Secretariat for Education, Research and Innovation (SERI) (M822.00025, SERI-funded ERC Starting Grant) and by funds from the Kanton Basel-Stadt and Basel-Land. We thank Alexander Schier, Everton Agnes, as well as the entire Kempf lab for valuable feedback on the manuscript.

## Author contributions

Conceptualization: G.A., S.T.S, M.A. and A.K. Investigation: S.T.S., G.A., C.K., L.M., M. A. V. Writing - original draft: S.T.S., Writing - Review and Editing: S.T.S., G.A., M. A. V., M.A., C.K., A.K. Funding acquisition: M.A. and A.K.

## Statistical analysis

Statistical analyses were performed in GraphPad Prism using the Mann-Whitney test.

