## Supplementary Material for "Tag-stabilized fluorescent protein binders for endogenous cell type-specific protein labelling"

Schnider Sophie Theodora *et al.*

### corresponding authors:

This PDF file includes:

Figures S1 to S6
Tables S1 to S4

#### Supplementary Figures and Figure Legends

Figure S1

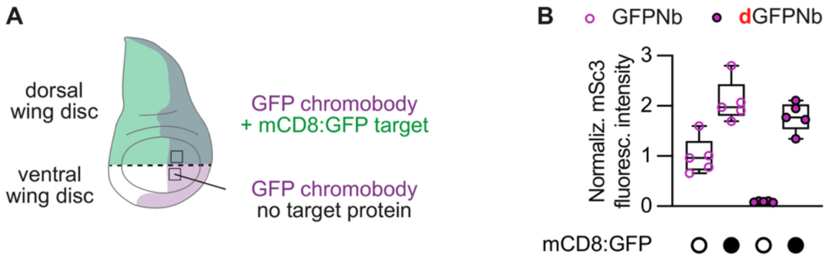

**Figure S1. Experimental setup and fluorescence measurements underlying GFP chromobody characterization. (A)** Schematic representation of the experimental strategy showing the wing imaginal disc with the target protein specifically expressed in the dorsal but not ventral compartment (using *ap*-LexA > *LexAop*-mCD8:GFP). The GFP chromobody was expressed in the posterior compartment using the *hh*-Gal4 genetic driver line (*hh*-Gal4 > *UAS*-GFPNb:mSc3 or *hh*-Gal4 > *UAS*-dGFPNb:mSc3). Squares indicate the regions that were analysed. **(B)** Quantification of mCD8:GFP-dependent mSc3 fluorescence intensities of GFPNb:mSc3 and dGFPNb:mSc3. Fluorescence values were normalized to the mean intensity of the GFPNb:mSc3 condition in the absence of the mCD8:GFP target protein. Data are shown as box-and-whisker plots (min to max) with all points displayed, $N = 5$. $N$, number of technical replicates. For detailed genotype descriptions, see Table S4.

Figure S2

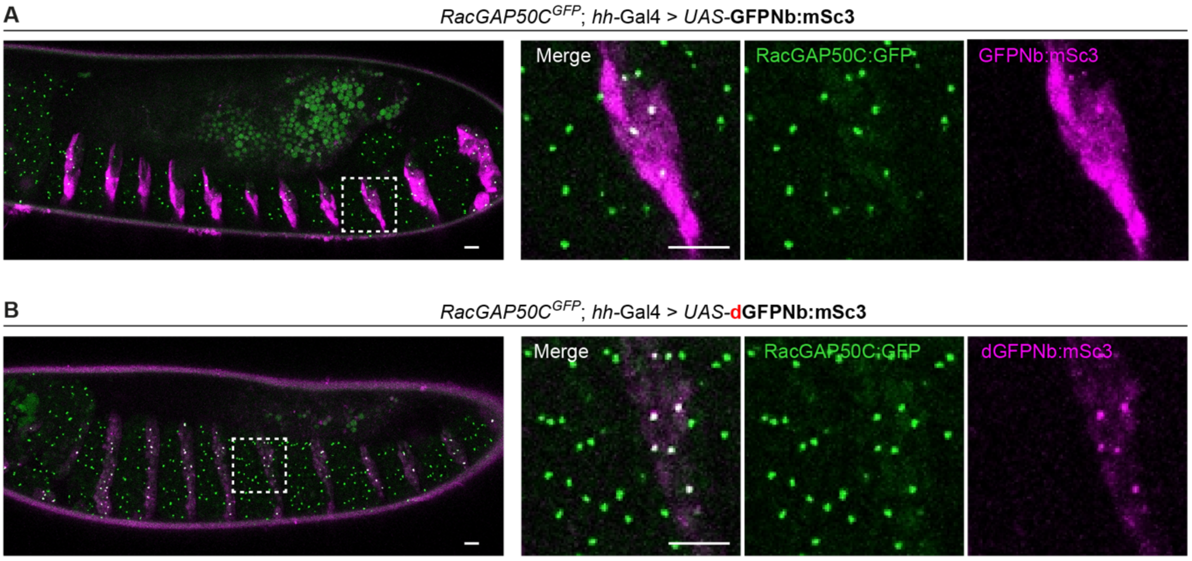

**Figure S2. Labelling endogenously-tagged RacGAP50C:GFP with GFP chromobodies.** Confocal live microscopy images of stage 14-15 embryos carrying a *RacGAP50C$^{GFP}$* knock-in allele and expressing either GFPNb:mSc3 **(A)** or dGFPNb:mSc3 **(B)** in posterior-compartment epidermal cells using *hh*-Gal4. Magnified views show epidermal cells spanning the boundary between anterior and posterior segmental compartments. Scale bars: 10 µm. For detailed genotype descriptions, see Table S4.

Figure S3

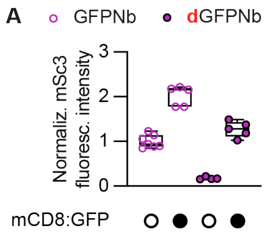

**Figure S3. Quantification of target-dependent GFP chromobody fluorescence in dFB neurons. (A)** GFPNb:mSc3 and dGFPNb:mSc3 were expressed in dFB neurons using *R23E10*-Gal4, with or without the mCD8:GFP target. Representative images are shown in Figure 3B and 3C. Fluorescence values measured in replicate ROIs were normalized to the mean intensity of the GFPNb:mSc3 condition in the absence of the mCD8:GFP target protein. Data are shown as box-and-whisker plots (min to max) with all points displayed, *N* = 4-7. *N*, number of technical replicates. For detailed genotype descriptions, see Table S4.

Figure S4

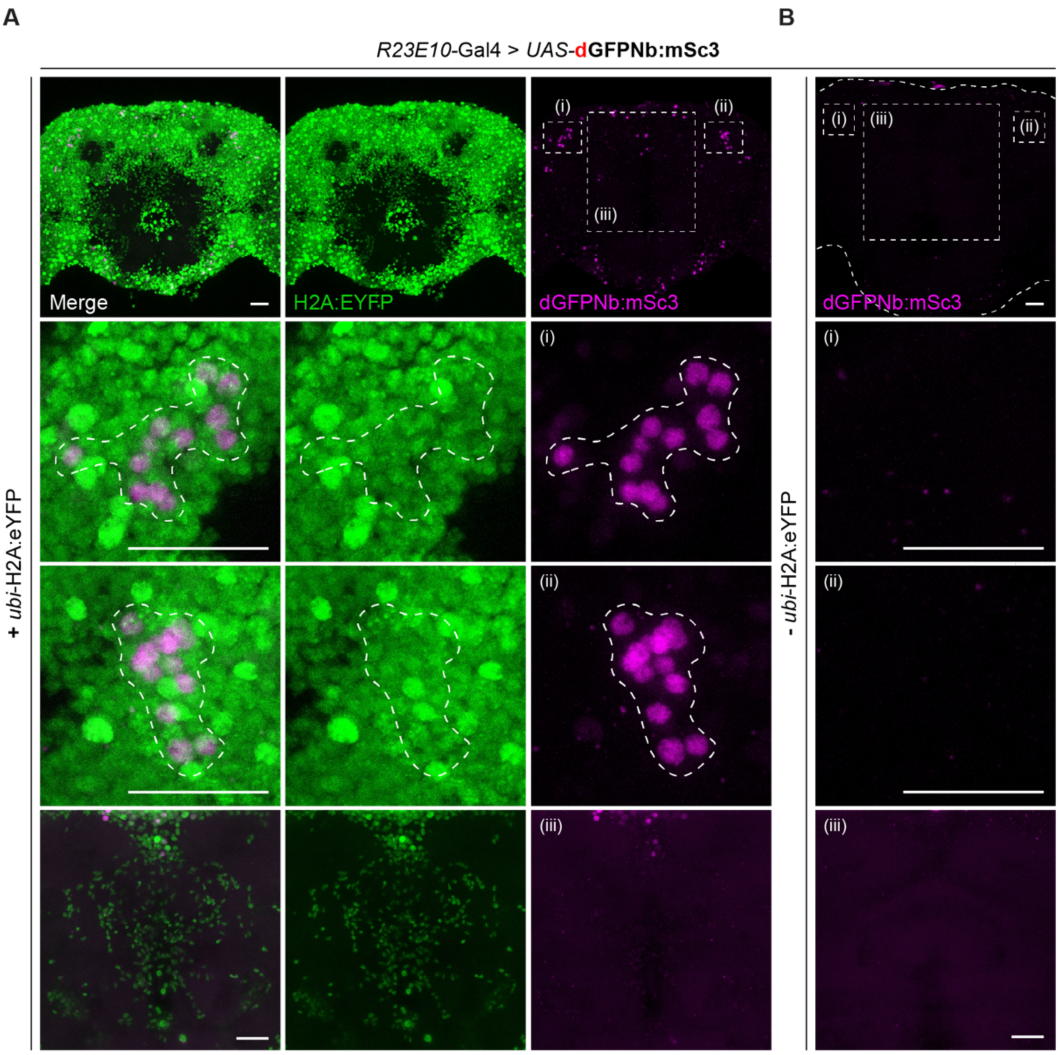

**Figure S4. The destabilized GFP chromobody labels nuclear H2A:eYFP in dFB neurons.** **(A and B)** Confocal microscopy images of adult brains in the presence **(A)** or absence **(B)** of *ubi*-H2A:eYFP and expressing dGFPNb:mSc3 in dFB neurons using *R23E10*-Gal4. Insets show magnified views of the left (i) and right (ii) groups of cell bodies, as well as of the fan-shaped body (iii). Dotted lines outline cell bodies of dorsal fan-shaped body neurons. Scale bars: 25 µm. For detailed genotype descriptions, see Table S4.

**Figure S5**

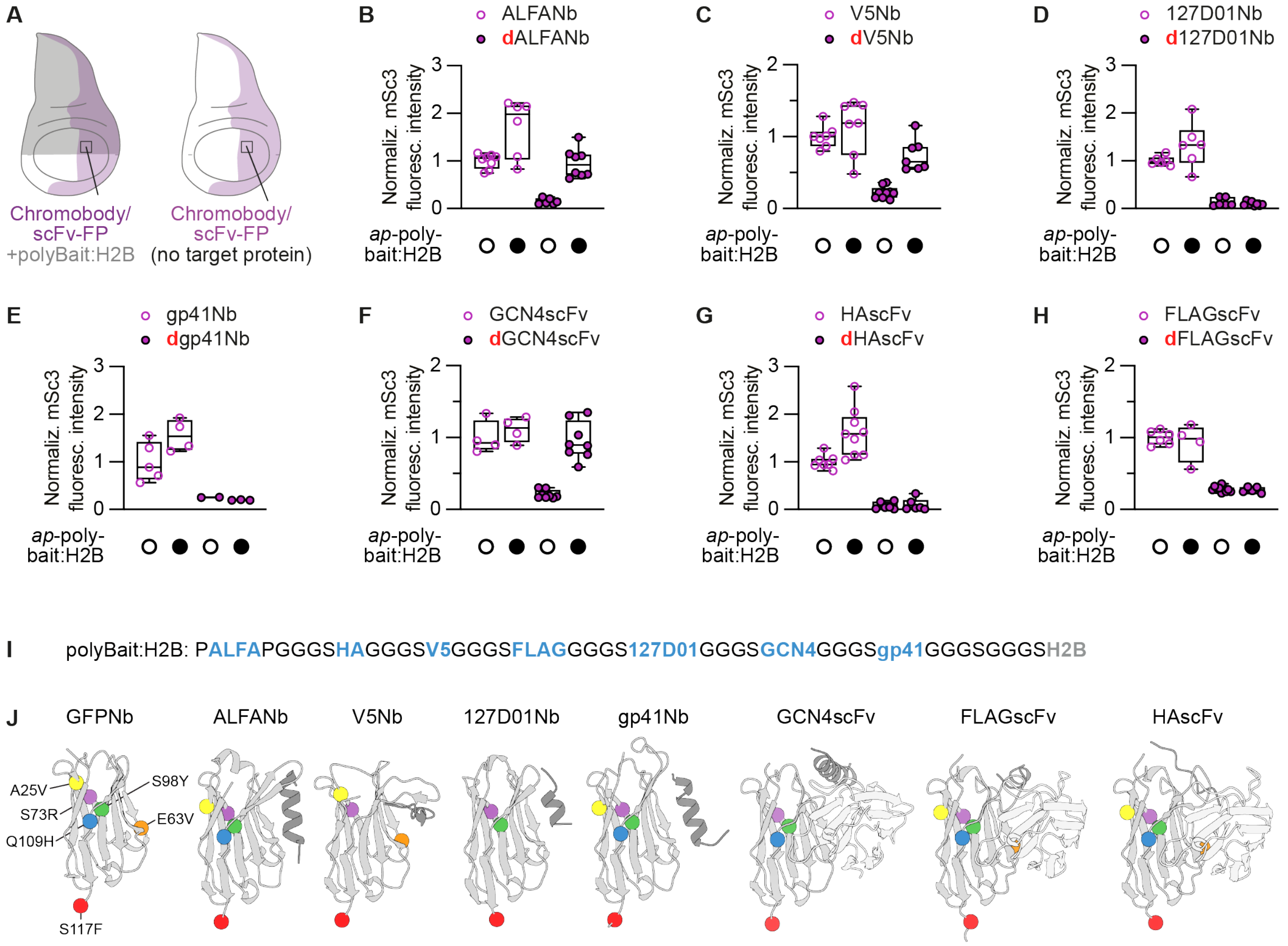

**Figure S5. Experimental design, target-dependent fluorescence, and structural analysis of short epitope tag chromobodies and scFvs. (A)** Schematic representation of the experimental strategy showing the wing imaginal disc with the target protein expressed in the dorsal but not ventral compartment (using *ap*-polyBait:H2B). The chromobody/scFv-FP fusion was expressed in the posterior compartment using the *hh*-Gal4 genetic driver line (*hh*-Gal4> *UAS*-chromobody/scFv-FP). Squares indicate the regions that were analysed. **(B-H)** Quantifications of polyBait:H2B-dependent mSc3 fluorescence intensities of ALFANb:mSc3 and dALFANb:mSc3 (B), V5Nb:mSc3 and dV5Nb:mSc3 (C), 127D01Nb:mSc3 and d127D01Nb:mSc3 (D), gp41Nb:mSc3 and dgp41Nb:mSc3 (E), GCN4scFv:mSc3 and dGCN4scFv:mSc3 (F), HAscFv:mSc3 and dHAscFv:mSc3 (G) and FLAGscFv:mSc3 and dFLAGscFv:mSc3 (H). Fluorescence values were normalized to the mean intensity of the corresponding stable chromobody or scFv:mSc3 fusion in the absence of polyBait:H2B target protein. **(I)** Amino acid sequence of the polyBait:H2B fusion. Short tags (blue) and linkers (grey) are indicated. **(J)** Structures or structural predictions of the tag-binding nanobodies and scFvs: GFP nanobody (PDB: 3OGO), ALFA nanobody with ALFA peptide (PDB: 6I2G), V5 nanobody with V5 peptide (PDB: 8SKJ), 127D01 nanobody with 127D01 peptide (AlphaFold 3 model), gp41 nanobody with gp41 peptide (AlphaFold 3 model), GCN4 scFv with GCN4 peptide (AlphaFold 3 model), HA scFv with HA peptide (AlphaFold 3 model), FLAG scFv with FLAG peptide (AlphaFold 3 model). Coloured spheres indicate the positions of the destabilizing substitutions. Data are shown as box-and-whisker plots (min to max) with all points displayed, *N* = 2-10. *N*, number of technical replicates. For detailed genotype descriptions, see Table S4.

Figure S6

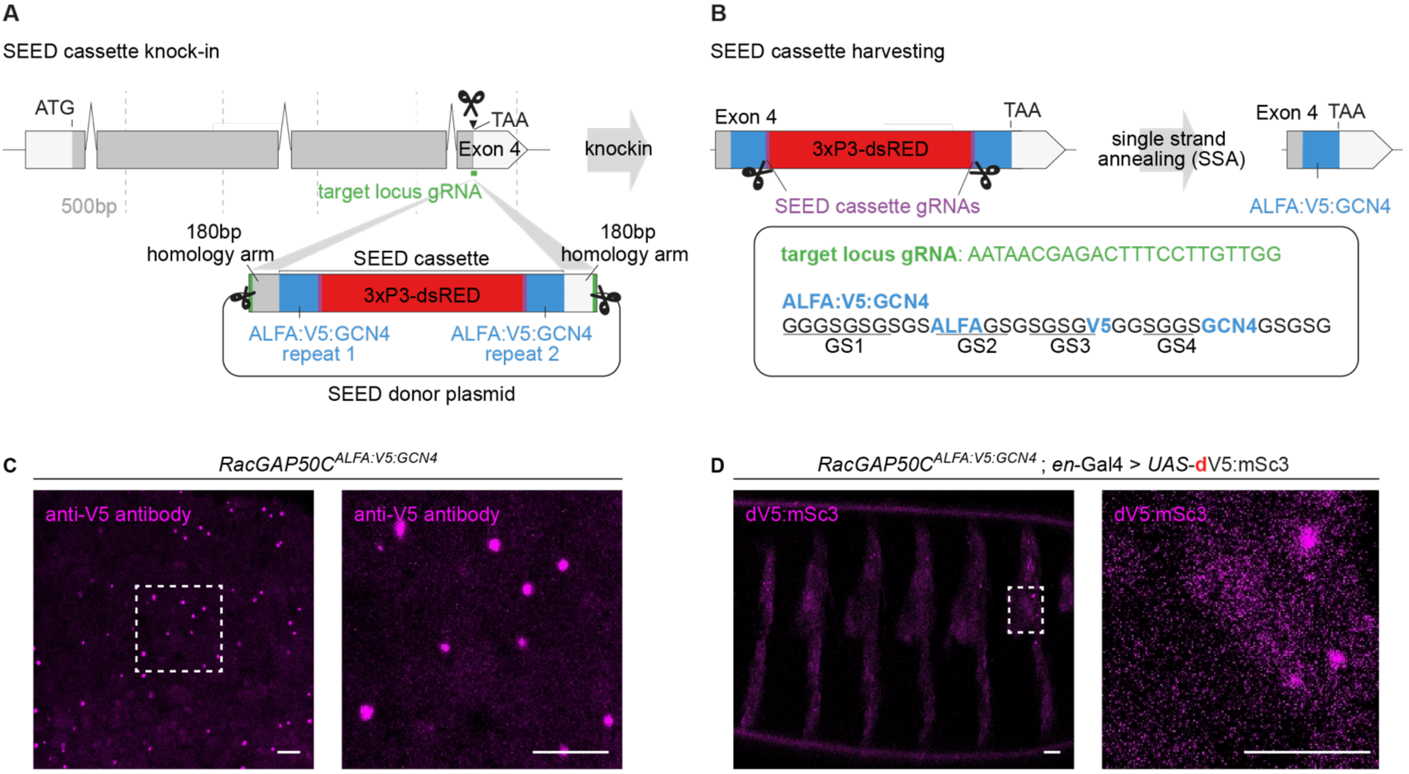

**Figure S6. SEED/Harvest generation and characterization of epitope-tagged RacGAP50C. (A and B)** Schematics of *RacGAP50C*$^{ALFA:V5:GCN4}$ knock-in generation (A) and harvesting strategy (B) using SEED/Harvest technology (see materials and methods). **(C)** Confocal microscopy image of epidermal cells from a stage 11 drosophila embryo carrying the *RacGAP50C*$^{ALFA:V5:GCN4}$ knock-in allele and stained with an anti-V5 antibody. **(D)** Confocal microscopy image of a stage 17 drosophila embryo carrying the *RacGAP50C*$^{ALFA:V5:GCN4}$ knock-in allele and expressing dV5Nb:mSc3 in posterior-compartment epidermal cells under the control of *en*-Gal4. Magnified views show epidermal cells spanning the boundary between anterior and posterior segmental compartments. Scale bars: 10 µm. For detailed genotype descriptions, see Table S4.

**Table S1.**
Amino acid sequences of nanobodies, scFvs and fluorescent proteins.

| |
|---|
| **GBP1**<br>MADVQLVESGGALVQPGGSLRLSCAASGFPVNRYSMRWYRQAPGKEREWVAGMSSA<br>GDRSSYEDSVKGRFTISRDDARNTVYLQMNSLKPEDTAVYYSNVNVGFEYWGQGTQVT<br>VSS |
| **dGBP1**<br>MADVQLVESGGALVQPGGSLRLSCVASGFPVNRYSMRWYRQAPGKEREWVAGMSSA<br>GDRSSYVDSVKGRFTIRRDDARNTVYLQMNSLKPEDTAVYYYNVNVGFEYWGHGTQVT<br>VSF |
| **ALFANb**<br>SGEVQLQESGGGLVQPGGSLRLSCTASGVTISALNAMAMGWYRQAPGERRVMVAAVS<br>ERGNAMYRESVQGRFTVTRDFTNKMVSLQMDNLKPEDTAVYYCHVLEDRVDSFHDYW<br>GQGTQVTVSS |
| **dALFANb**<br>MSGEVQLQESGGGLVQPGGSLRLSCVASGVTISALNAMAMGWYRQAPGERRVMVAAV<br>SERGNAMYRESVQGRFTVRRDFTNKMVSLQMDNLKPEDTAVYYYHVLEDRVDSFHDY<br>WGHGTQVTVSF |
| **V5Nb**<br>MAEVQLQASGGGFVQPGGSLRLSCAASGTTSFGDTMGWFRQAPGKEREFVSAISRQG<br>DKHYYADSVKGRFTISRDNSKNTVYLQMNSLRAEDTATYYCAEWMNTRREFITPYWGQ<br>GTQVTVSS |
| **dV5Nb$^{4mut}$**<br>MAEVQLQASGGGFVQPGGSLRLSCVASGTTSFGDTMGWFRQAPGKEREFVSAISRQG<br>DKHYYVDSVKGRFTIRRDNSKNTVYLQMNSLRAEDTATYYCAEWMNTRREFITPYWGQ<br>GTQVTVSF |
| **MoontagNb**<br>MEVQLVESGGGLVQPGGSLRLSCAASGSISSVDVMSWYRQAPGKQRELVAFITDRGRT<br>NYKVSVKGRFTISRDNSKNMVYLQMNSLKPEDTADYLCRAESRTSWSSPSPLDVWGRG<br>TQVTVSSL |
| **dMoontagNb**<br>MEVQLVESGGGLVQPGGSLRLSCVASGSISSVDVMSWYRQAPGKQRELVAFITDRGRT<br>NYKVSVKGRFTIRRDNSKNMVYLQMNSLKPEDTADYLYRAESRTSWSSPSPLDVWGHG<br>TQVTVSFL |
| **127D01Nb**<br>MEVQLVESGGGLVQAGESLRLSCAASGSTFDFKVMGWYRQPPGKQREGVAAIRLSGN<br>MHYAESVKGRFAISKANAKNTVYLQMNSLRPEDTAVYYCKVNIRGQDYWGQGTQVTVS<br>S |
| **d127D01Nb**<br>MEVQLVESGGGLVQAGESLRLSCAASGSTFDFKVMGWYRQPPGKQREGVAAIRLSGN<br>MHYAESVKGRFAIRKANAKNTVYLQMNSLRPEDTAVYYYKVNIRGQDYWGQGTQVTVS<br>F |
| **GCN4scFv**<br>MGPDIVMTQSPSSLSASVGDRVTITCRSSTGAVTTSNYASWVQEKPGKLFKGLIGGTNN<br>RAPGVPSRFSGSLIGDKATLTISSLQPEDFATYFCALWYSNHWVFGQGTKVELKRGGGG<br>SGGGGSGGGGSGGGGSEVKLLESGGGLVQPGGSLKLSCAVSGFSLTDYGVNWVRQA<br>PGRGLEWIGVIWGDGITDYNSALKDRFIISKDNGKNTVYLQMSKVRSDDTALYYCVTGLF<br>DYWGQGTLVTVSS |
| **dGCN4scFv**<br>MGPDIVMTQSPSSLSASVGDRVTITCRSSTGAVTTSNYASWVQEKPGKLFKGLIGGTNN<br>RAPGVPSRFSGSLIGDKATLTISSLQPEDFATYFCALWYSNHWVFGQGTKVELKRGGGG |

| |
|---|
| SGGGGSGGGGSGGGGSEVKLLESGGGLVQPGGSLKLSCAVSGFSLTDYGVNWVRQA<br>PGRGLEWIGVIWGDGITDYNSALKDRFIIRKDNGKNTVYLQMSKVRSDDTALYYYVTGLF<br>DYWGHGTLVTVSF |
| **HAscFv**<br>MAEVKLVESGGGLVKPGGSLKLSCAASGFTFSSYGMSWVRQTPEKRLEWVATISRGGS<br>YTYYPDSVKGRFTISRDNAKNTLYLQMSSLRSEDTAIYYCARRETYDEKGFAYWGQGTT<br>LTVSSGGGGSGGGGSGGGGSDIVLTQSPASLTVSLGQRATISCKSSQSLLNSGNQKNYL<br>TWYQQKPGQPPKLLIYWASTRESGIPARFSGSGSGTDFTLNIHPVEEEDAATYYCQNDN<br>SHPLTFGAGTKLEIKRAAAKG |
| **dHAscFv**<br>MAEVKLVESGGGLVKPGGSLKLSCVASGFTFSSYGMSWVRQTPEKRLEWVATISRGGS<br>YTYYVDSVKGRFTIRRDNAKNTLYLQMSSLRSEDTAIYYYARRETYDEKGFAYWGHGTT<br>LTVSFGGGGSGGGGSGGGGSDIVLTQSPASLTVSLGQRATISCKSSQSLLNSGNQKNYL<br>TWYQQKPGQPPKLLIYWASTRESGIPARFSGSGSGTDFTLNIHPVEEEDAATYYCQNDN<br>SHPLTFGAGTKLEIKRAAAKG |
| **FLAGscFv**<br>MAEVKLVESGGGLVKPGGSLKLSCAASGFTFSSFGMHWVRQTPEKRLEWVAYISSGSS<br>TIYYADTVKGRFTISRDNAKNTLYLQMSSLRSEDTAIYYCARSLATAAFAYWGQGTTLTVS<br>SGGGGSGGGGSGGGGSDIVLTQSPASLTVSLGQRATISCRSSQSIVYSNGNTYLEWYQ<br>QKPGQPPKLLIYKVSNRFSGIPARFSGSGSGTDFTLNIHPVEEEDAATYYCFQGSHVPYT<br>FGAGTKLEIKRAAAKG |
| **dFLAGscFv**<br>MAEVKLVESGGGLVKPGGSLKLSCVASGFTFSSFGMHWVRQTPEKRLEWVAYISSGSS<br>TIYYVDTVKGRFTIRRDNAKNTLYLQMSSLRSEDTAIYYYARSLATAAFAYWGHGTTLTVS<br>FGGGGSGGGGSGGGGSDIVLTQSPASLTVSLGQRATISCRSSQSIVYSNGNTYLEWYQ<br>QKPGQPPKLLIYKVSNRFSGIPARFSGSGSGTDFTLNIHPVEEEDAATYYCFQGSHVPYT<br>FGAGTKLEIKRAAAKG |
| **mScarlet3 (mSc3)**<br>MDSTEAVIKEFMRFKVHMEGSMNGHEFEIEGEGEGRPYEGTQTAKLRVTKGGPLPFSW<br>DILSPQFMYGSRAFTKHPADIPDYWKQSFPEGFKWERVMNFEDGGAVSVAQDTSLEDG<br>TLIYKVKLRGTNFPPDGPVMQKKTMGWEASTERLYPEDVVLKGDIKMALRLKDGGRYLA<br>DFKTTYRAKKPVQMPGAFNIDRKLDITSHNEDYTVVEQYERSVARHSTGGSGGS |
| **mGreenLantern (mGL)**<br>MVSKGEELFTGVVPILVELDGDVNGHKFSVRGEGEGDATNGKLTLKFICTTGKLPVPWPT<br>LVTTLGYGVACFARYPDHMKQHDFFKSAMPEGYVQERTISFKDDGTYKTRAEVKFEGDT<br>LVNRIVLKGIDFKEDGNILGHKLEYNFNSHKVYITADKQKNGIKANFKTRHNVEDGGVQLA<br>DHYQQNTPIGDGPVLLPDNHYLSHQSKLSKDPNEKRDHMVLKERVTAAGITHDMDELYK |
| **Tdx5StayGold (tdSG)**<br>MVSTGEELFTGVVPFKFQLKGTINGKSFTVEGEGEGNSHEGSHKGKYVCTSGKLPMSW<br>AALGTSFGYGMKYYTKYPSGLKNWFHEVMPEGFTYDRHIQYKGDGSIHAKHQHFMKNG<br>TYHNIVEFTGQDFKENSPVLTGDMNVSLPNEVQHIPRDDGVECPVTLLYPLLSDKSKCVE<br>AYQNTIIKPLHNQPAPDVPYHWIRKQYTQSKDDTEERDHIIQSETLEAHLPWHEPSASAV<br>EFGHGTGSTGSGSSGTASSEDNNKLMVSTGEELFTGVVPFKFQLKGTINGKSFTVEGEG<br>EGNSHEGSHKGKYVCTSGKLPMSWAALGTSFGYGMKYYTKYPSGLKNWFHEVMPEGF<br>TYDRHIQYKGDGSIHAKHQHFMKNGTYHNIVEFTGQDFKENSPVLTGDMNVSLPNEVQH<br>IPRDDGVECPVTLLYPLLSDKSKCVEAYQNTIIKPLHNQPAPDVPYHWIRKQYTQSKDDTE<br>ERDHIIQSETLEAHL |
| **mTagBFP2**<br>MVSKGEELIKENMHMKLYMEGTVDNHHFKCTSEGEGKPYEGTQTMRIKVVEGGPLPFA<br>FDILATSFLYGSKTFINHTQGIPDFFKQSFPEGFTWERVTTYEDGGVLTATQDTSLQDGCL<br>IYNVKIRGVNFTSNGPVMQKKTLGWEAFTETLYPADGGLEGRNDMALKLVGGSHLIANA<br>KTTYRSKKPAKNLKMPGVYYVDYRLERIKEANNETYVEQHEVAVARYCDLPSKLGHKLN |

| |
|---|
| **dendra2**<br>MNTPGINLIKEDMRVKVHMEGNVNGHAFVIEGEGKGKPYEGTQTANLTVKEGAPLPFSY<br>DILTTAVHYGNRVFTKYPEDIPDYFKQSFPEGYSWERTMTFEDKGICTIRSDISLEGDCFF<br>QNVRFKGTNFPPNGPVMQKKTLKWEPSTEKLHVRDGLLVGNINMALLLEGGGHYLCDF<br>KTTYKAKKVVQLPDAHFVDHRIEILGNDSDYNKVKLYEHAVARYSPLPSQVW |
| **polyBait**<br>MPSRLEEELRRRLTEPGGGSYPYDVPDYAGGGSGKPIPNPLLGLDSTGGGSDYKDDDD<br>KGGGSSFEDFWKGEDGGGSEELLSKNYHLENEVARLKKGGGSKNEQELLELDKWASL<br>GGGSGGGSMPPKTSGKAAKKAGKAQKNITKTDKKKKRKRKESYAIYIYKVLKQVHPDTGI<br>SSKAMSIMNSFVNDIFERIAAEASRLAHYNKRSTITSREIQTAVRLLLPGELAKHAVSEGTK<br>AVTKYTSSK |

**Table S2.**
Mutations in nanobodies and scFvs.

| Protein binder | AA | sequence context | mutate to | number | position structure | annotation | Tang et al. |
|---|---|---|---|---|---|---|---|
| GFPNb | A | LSCAASG | V | 25 | **3OGO**<br>25 | A25V | A25V |
| | E | SSYEDSV | V | 63 | 62 | E63V | E68V |
| | S | FTISRDD | R | 73 | 72 | S73R | S79R |
| | S | VYYSNVN | Y | 98 | 97 | S98Y | S104Y |
| | Q | YWGQGTQ | H | 109 | 108 | Q109H | Q120H |
| | S | TVSS | F | 117 | 116 | S117F | S128F |
| ALFANb | T | LSCTASG | V | 26 | **6i2g**<br>25 | T26V | |
| | R | AMYRESV | | | | | |
| | T | FTVTRDF | R | 76 | 75 | T76R | |
| | C | VYYCHVL | Y | 101 | 100 | C101Y | |
| | Q | YWGQGTQ | H | 117 | 116 | Q117H | |
| | S | TVSS | F | 125 | 124 | S125F | |
| V5Nb | A | LSCAASG | V | 25 | **8SKJ**<br>26 | A25V | |
| | A | HYYADSV | V | 63 | 64 | A63V | |
| | S | FTISRDN | R | 73 | 74 | S73R | |
| | C | TYYCAEW | | | | | |
| | Q | YWGQGTQ | | | | | |
| | S | TVSS | F | 123 | 124 | S123F | |
| 127D01Nb | A | LSCAASG | | | **Alphafold3** | | |
| | A | MHYAESV | | | | | |
| | S | FAISKAN | R | 71 | 71 | S71R | |
| | C | VYYCKVN | Y | 96 | 96 | C96Y | |
| | Q | YWGQGTQ | | | | | |
| | S | TVSS | F | 116 | 116 | S116F | |
| gp41Nb | A | SCAASGS | V | 24 | **7AEJ, Alphafold3 M**<br>24, 25 | A24V | |
| | K | TNYKVSV | | | | | |
| | S | FTISRDN | R | 71 | 70, 71 | S71R | |
| | **C** | DYLCRAE | Y | 96 | 92, 96 | C96Y | |
| | R | VWGRGTQ | H | 115 | 105, 115 | R115H | |
| | S | TVSSL-- | F | 123 | N/A, 123 | S123F | |
| GCN4scFv VH | A | LSCAVSG | | | **1P4I, Alphafold3 M** | | |
| | N | TDYNSAL | | | | | |
| | S | FIISKDN | R | 73 | VH T81, 204 | S73R | |

| | | | | | | |
|---|---|---|---|---|---|---|
| | C | LYYCVTG | Y | 98 | VH 106,229 | C98Y |
| | Q | YWGQGTL | H | 108 | VH 141,239 | Q108H |
| | S | TVSS | F | 116 | VH 148, 247 | S116F |
| HAscFv VH | A | LSCAASG | V | 25 | **Alphafold3 M** 25 | A25V |
| | P | TYYPDSV | V | 63 | 63 | P63V |
| | S | FTISRDN | R | 73 | 73 | S73R |
| | C | IYYCARR | Y | 98 | 98 | C98Y |
| | Q | YWGQGTT | H | 114 | 114 | Q114H |
| FLAGscFv VH | A | LSCAASG | V | 25 | **Alphafold3 M** 25 | A25V |
| | A | IYYADTV | V | 63 | 63 | A63V |
| | S | FTISRDN | R | 73 | 73 | S73R |
| | C | IYYCARS | Y | 98 | 98 | C98Y |
| | Q | YWGQGTT | H | 112 | 112 | Q112H |
| | S | TVSS | F | 120 | 120 | S120F |

**Table S3.**
DNA constructs for RacGAP50C$^{ALFA:V5:GCN4}$ based on the SEED/harvest technology.

| Construct | Sequence |
|---|---|
| **gRNA** | AATAACGAGACTTTCCTTGTTGG |
| **180bp homology arm 5'** | CAGTTATCAAGCGGGTGCCAAGCAACAAAAATGACTTACTCTC<br>GCTATATGGTGAGTTATTTATGAGCTAATTTTTTTAATTATTCTAA<br>TTATCTCTCCTTCACAGCAACTCCCTTCAAAGGAGGCACCATTA<br>AGAAGCGGAAGTTTTACGGCACACCGCCGGCATCTGCGCACA<br>AGAAA |
| **180bp homology arm 3'** | TAACGAGACTTTCCTTGTTGTAAAGCATTTGACGTGTTACTTAA<br>GTTTACTTCAAGTATTTTGCAAGTTTTTTGTTTCCGGCCAGCCAA<br>CCCCGAGCTGTATTTGTACCCAACCCAACCATAAGCTTTTCTTT<br>ATAATTTATGCGTTCGATTATAAACTAAAGCACGTTATTATCGTA<br>CT |
| **SEED-ALFA:V5:GCN4** | GGAGGCGGAAGCGGCTCTGGCTCTGGAtcaCGTTTGGAAGAGG<br>AACTCAGACGCCGCTTaACTGaaGGGTCTGGATCTGGATCAGG<br>TAAGCCTATCCCTAACCCTCTCCTCGGTCTTGATTCTACGGGTG<br>GGTCGGGCGGCTCCGAAGAACTTTTGAGCAAGAATTATCATCT<br>TGAGAACGAAGTGGCTCGTCTTAAGAAAGGATCCTacCCTAGGt<br>agCCGCGTTGTTATGATCGTACTACGCTAGCGTAGTACGATCAT<br>AACAACGCGGGTAGTACGATCATAACAACGCGGGGATCTAATT<br>CAATTAGAGACTAATTCAATTAGAGCTAATTCAATTAGGATCCAA<br>GCTTATCGATTTCGAACCCTCGACCGCCGGAGTATAAATAGAG<br>GCGCTTCGTCTACGGAGCGACAATTCAATTCAAACAAGCAAAG<br>TGAACACGTCGCTAAGCGAAAGCTAAGCAAATAAACAAGCGCA<br>GCTGAACAAGCTAAACAATCGGCTCGAAGCCGGTCGCCACCatg<br>GCCTCCTCCGAGGACGTCATCAAGGAGTTCATGCGCTTCAAGG<br>TGCGCATGGAGGGCTCCGTGAACGGCCACGAGTTCGAGATCG<br>AGGGCGAGGGCGAGGGCCGCCCCTACGAGGGCACCCAGACC<br>GCCAAGCTGAAGGTGACCAAGGGCGGCCCCCTGCCCTTCGCC<br>TGGGACATCCTGTCCCCCCAGTTCCAGTACGGCTCCAAGGTGT<br>ACGTGAAGCACCCCGCCGACATCCCCGACTACAAGAAGCTGTC<br>CTTCCCCGAGGGCTTCAAGTGGGAGCGCGTGATGAACTTCGA<br>GGACGGCGGCGTGGTGACCGTGACCCAGGACTCCTCCCTcCA<br>GGACGGCTCCTTCATCTACAAGGTGAAGTTCATCGGCGTGAAC<br>TTCCCCTCCGACGGCCCCGTAATGCAGAAGAAGACTATGGGCT<br>GGGAGGCgTCCACCGAGCGCCTGTACCCCCGCGACGGCGTGC<br>TGAAGGGCGAGATCCACAAGGCCCTGAAGCTGAAGGACGGCG<br>GCCACTACCTGGTGGAGTTCAAGTCCATCTACATGGCCAAGAA<br>GCCCGTGCAGCTGCCCGGCTACTACTACGTGGACTCCAAGCT<br>GGACATCACCTCCCACAACGAGGACTACACCATCGTGGAGCAG<br>TACGAGCGCGCCGAGGGCCGCCACCACCTGTTCCTGTAGCGG<br>CCGCGACTCTAGATCATAATCAGCCATACCACATTTGTAGAGGT<br>TTTACTTGCTTTAAAAAACCTCCCACACCTCCCCCTGAACCTGA<br>AACATAAAATGAATGCAATTGTTGTTGTTAACTTGTTTATTGCAG<br>CTTATAATGGTTACAAATAAAGCAATAGCATCACAAATTTCACAA<br>ATAAAGCATTTTTTTCACTGCATTCTAGTTGTGGTTTGTCCAAAC<br>TCATCAATGTATCTTACCGCTGGTACTGTATGGATGTACCCGCT<br>GGTACTGTATGGATGTACGCTAGCGTACATCCATACAGTACCA<br>GCGGACTAGTAGGCTCTGGAtcaCGTTTGGAAGAGGAACTCAGA<br>CGCCGCTTaACTGaaGGGTCTGGATCTGGATCAGGTAAGCCTA |

| | |
|---|---|
| | TCCCTAACCCTCTCCTCGGTCTTGATTCTACGGGTGGGTCGGGCGGCTCCGAAGAACTTTTGAGCAAGAATTATCATCTTGAGAACGAAGTGGCTCGTCTTAAGAAAGGATCCGGtTCCGGC<br>Linker<br>Tag (ALFA, V5, GCN4)<br>SEED guide RNAs<br>3xP3-Hsp70promoter<br>dsRed |

**Table S4.**
Detailed description of genotypes per Figure.

| Figure | Name in Figure | Genotype |
|---|---|---|
| Figure 1C | *ap*-LexA > *LexAop*-mCD8:GFP;*hh*-Gal4 > *UAS*-GFPNb:mSc3 | *;ap-LexA, LexAop-mCD8:GFP/CyO;hh-Gal4/UAS-GFPNb:Sc3* |
| Figure 1D | *ap*-LexA > *Lex:Aop*-mCD8:GFP;*hh*-Gal4 > *UAS*-dGFPNb:mSc3 | *;ap-LexA, LexAop-mCD8:GFP/CyO;hh-Gal4/UAS-dGFPNb:Sc3* |
| Figure 1E | GFPNb:mSc3 | *;ap-LexA, LexAop-mCD8:GFP/CyO;hh-Gal4/UAS-GFPNb:Sc3* |
| Figure 1E | dGFPNb:mSc3 | *;ap-LexA, LexAop-mCD8:GFP/CyO;hh-Gal4/UAS-dGFPNb:Sc3* |
| Figure 2A,B upper panel | $Ecad^{GFP}$, *btl*-Gal4 > *UAS*-GFPNb:mSc3 | ;$Ecad^{GFP}$ */+;btl-Gal4/UAS-GFPNb:Sc3* |
| Figure 2A,B lower panel | $Ecad^{GFP}$, *btl*-Gal4 > *UAS*-dGFPNb:mSc3 | ;$Ecad^{GFP}$ */+;btl-Gal4/UAS-dGFPNb:Sc3* |
| Figure 2C,D upper panel | $ZO\text{-}1^{eGFP}$; *fli1a*-GFPNb:mSc3 | $ZO\text{-}1^{eGFP}$; *fli1a*-GFPNb:mSc3 |
| Figure 2C,D lower panel | $ZO\text{-}1^{eGFP}$; *fli1a*-dGFPNb:mSc3 | $ZO\text{-}1^{eGFP}$; *fli1a*-dGFPNb:mSc3 |
| Figure 3B | *R23E10*-Gal4 > *UAS*-GFPNb:mSc3 +mCD8:GFP | *;UAS-mCD8:GFP/CyO;R23E10-Gal4/UAS-GFPNb:mSc3* |
| Figure 3C | *R23E10*-Gal4 > *UAS*-dGFPNb:mSc3 +mCD8:GFP | *;UAS-mCD8:GFP/CyO;R23E10-Gal4/UAS-GFPNb:mSc3* |
| Figure 3D | GFPNb:mSc3 | *;UAS-mCD8:GFP/CyO;R23E10-Gal4/UAS-GFPNb:mSc3* |
| Figure 3D | dGFPNb:mSc3 | *;UAS-mCD8:GFP/CyO;R23E10-Gal4/UAS-dGFPNb:mSc3* |
| Figure 3E left panels | *R23E10*-Gal4 > *UAS*-dGFPNb:mSc3 +Cv-$c^{GFP}$ | *;if/CyO; Cv*-$c^{GFP}$*, R23E10-Gal4, UAS-dGFPNb:mSc3* |
| Figure 3E right panels | *R23E10*-Gal4 > *UAS*-dGFPNb:mSc3 | *;+/CyO; R23E10-Gal4, UAS-dGFPNb:mSc3* |
| Figure 4C | *hh*-Gal4 > *UAS*-ALFANb | *;+/if;hh-Gal4/UAS-ALFANb:mSc3* |
| Figure 4C | *hh*-Gal4 > *UAS*-ALFANb+ *ap*-polyBait:H2B | *;+/ap-polyBait:H2B; hh-Gal4/UAS-ALFANb:mSc3* |

| | | |
|---|---|---|
| Figure 4C | *hh*-Gal4 > *UAS*-dALFANb | *;+/if;hh-Gal4/UAS-dALFANb:mSc3* |
| Figure 4C | *hh*-Gal4 > *UAS*-dALFANb+ *ap*-polyBait:H2B | *;+/ap-polyBait:H2B; hh-Gal4/UAS-dALFANb:mSc3* |
| Figure 4C | *hh*-Gal4 > *UAS*-V5Nb | *;+/if;hh-Gal4/UAS-V5Nb:mSc3* |
| Figure 4C | *hh*-Gal4 > *UAS*-V5Nb+*ap*-polyBait:H2B | *;+/ap-polyBait:H2B; hh-Gal4/UAS-V5Nb:mSc3* |
| Figure 4C | *hh*-Gal4 > *UAS*-dV5Nb | *;+/if;hh-Gal4/UAS-dV5Nb:mSc3* |
| Figure 4C | *hh*-Gal4 > *UAS*-V5Nb+*ap*-polyBait:H2B | *;+/ap-polyBait:H2B; hh-Gal4/UAS-dV5Nb:mSc3* |
| Figure 4C | *hh*-Gal4 > *UAS*-127D01Nb | *;+/if;hh-Gal4/UAS-127D01Nb:mSc3* |
| Figure 4C | *hh*-Gal4 > *UAS*-127D01Nb+ *ap*-polyBait:H2B | *;+/ap-polyBait:H2B; hh-Gal4/UAS-127D01Nb:mSc3* |
| Figure 4C | *hh*-Gal4 > *UAS*-d127D01Nb | *;+/if;hh-Gal4/UAS-d127D01Nb:mSc3* |
| Figure 4C | *hh*-Gal4 > *UAS*-d127D01Nb+*ap*-polyBait:H2B | *;+/ap-polyBait:H2B; hh-Gal4/UAS-d127D01Nb:mSc3* |
| Figure 4C | *hh*-Gal4 > *UAS*-gp41Nb | *;+/if;hh-Gal4/UAS-gp41Nb:mSc3* |
| Figure 4C | *hh*-Gal4 > *UAS*-gp41Nb+*ap*-polyBait:H2B | *;+/ap-polyBait:H2B; hh-Gal4/UAS-gp41Nb:mSc3* |
| Figure 4C | *hh*-Gal4 > *UAS*-dgp41Nb | *;+/if;hh-Gal4/UAS-dgp41Nb:mSc3* |
| Figure 4C | *hh*-Gal4 > *UAS*-dgp41Nb+*ap*-polyBait:H2B | *;+/ap-polyBait:H2B; hh-Gal4/UAS-dgp41Nb:mSc3* |
| Figure 4D | *hh*-Gal4 > *UAS*-GCN4scFv | *;+/if;hh-Gal4/UAS-GCN4scFv:mSc3* |
| Figure 4D | *hh*-Gal4 > *UAS*-GCN4scFv+*ap*-polyBait:H2B | *;+/ap-polyBait:H2B; hh-Gal4/UAS-GCN4scFv:mSc3* |
| Figure 4D | *hh*-Gal4 > *UAS*-dGCN4scFv | *;+/if;hh-Gal4/UAS-dGCN4scFv* |
| Figure 4D | *hh*-Gal4 > *UAS*-dGCN4scFv+*ap*-polyBait:H2B | *;+/ap-polyBait:H2B; hh-Gal4/UAS-dGCN4scFv* |
| Figure 4E | *hh*-Gal4 > *UAS*-HAscFv | *;+/if;hh-Gal4/UAS-HA:mSc3* |
| Figure 4E | *hh*-Gal4 > *UAS*-HAscFv+*ap*-polyBait:H2B | *;+/ap-polyBait:H2B; hh-Gal4/UAS-HA:mSc3* |
| Figure 4E | *hh*-Gal4 > *UAS*-dHAscFv | *;+/if;hh-Gal4/UAS-dHAscFv* |
| Figure 4E | *hh*-Gal4 > *UAS*-dHAscFv+*ap*-polyBait:H2B | *;+/ap-polyBait:H2B; hh-Gal4/UAS-dFLAGscFv* |
| Figure 4E | *hh*-Gal4 > *UAS*-FLAGscFv | *;+/if;hh-Gal4/UAS-FLAG:mSc3* |
| Figure 4E | *hh*-Gal4 > *UAS*-FLAGscFv+*ap*-polyBait:H2B | *;+/ap-polyBait:H2B; hh-Gal4/UAS-FLAG:mSc3* |
| Figure 4E | *hh*-Gal4 > *UAS*-dFLAGscFv | *;+/if;hh-Gal4/UAS-dFLAGscFv* |

| | | |
|---|---|---|
| Figure 4E | *hh*-Gal4 > *UAS*-dFLAGscFv+*ap*-polyBait:H2B | *;+/ap-polyBait:H2B; hh-Gal4/UAS-dFLAGscFv* |
| Figure 4F | See Figure 4 C, D, E | See Figure 4 C, D, E |
| Figure 5C | *RacGAP50C*$^{GFP}$ | *RacGAP50C*$^{GFP}$ |
| Figure 5D | *RacGAP50C*$^{ALFA\text{-}V5\text{-}GCN4}$, *en*-Gal4>UAS-dALFA:mGL | *;RacGAP50C*$^{ALFA\text{-}V5\text{-}GCN4}$*/en-Gal4; dALFA:mGL/+* |
| Figure 5D | *RacGAP50C*$^{ALFA\text{-}V5\text{-}GCN4}$, *en*-Gal4>UAS-dGCN4:mSc3 | *;RacGAP50C*$^{ALFA\text{-}V5\text{-}GCN4}$*/en-Gal4; dGCN4:mSc3/+* |
| Figure S6C | *RacGAP50C*$^{ALFA\text{-}V5\text{-}GCN4}$ | *;RacGAP50C*$^{ALFA\text{-}V5\text{-}GCN4}$ |
| Figure S6D | *RacGAP50C*$^{ALFA\text{-}V5\text{-}GCN4}$, *en*-Gal4>UAS-dV5:mSc3 | *;RacGAP50C*$^{ALFA\text{-}V5\text{-}GCN4}$*/en-Gal4; dV5:mSc3/+* |
| **Supplementary Figure** | **Name in Figure** | **Genotype** |
| Figure S1B | GFPNb:mSc3+mCD8:GFP | *;ap-LexA, LexAop-mCD8:GFP/CyO;hh-Gal4/UAS-GFPNb:Sc3* |
| Figure S1B | dGFPNb:mSc3+mCD8:GFP | *;ap-LexA, LexAop-mCD8:GFP/CyO;hh-Gal4/UAS-dGFPNb:Sc3* |
| Figure S2A | *RacGAP50C*$^{GFP}$, *hh*-Gal4 >*UAS*-GFPNb:mSc3 | *;RacGAP50C*$^{GFP}$*/CyO or if; hh-Gal4/UAS-GFPNb:mSc3* |
| Figure S2B | *RacGAP50C*$^{GFP}$, *hh*-Gal4 >*UAS*-dGFPNb:mSc3 | *;RacGAP50C*$^{GFP}$*/CyO or if; hh-Gal4/UAS-dGFPNb:mSc3* |
| Figure S3A | GFPNb:mSc3+mCD8:GFP | *;UAS-mCD8:GFP/CyO;R23E10-Gal4/UAS-GFPNb:mSc3* |
| Figure S3A | dGFPNb:mSc3+mCD8:GFP | *;UAS-mCD8:GFP/CyO;R23E10-Gal4/UAS-dGFPNb:mSc3* |
| Figure S4A | *R23E10*-Gal4 > *UAS*-dGFPNb:mSc3 + *ubi*-H2A:eYFP | *ubi-H2A:eYFP/+;if/CyO; R23E10-Gal4, UAS-dGFPNb:mSc3/+* |
| Figure S4B | *R23E10*-Gal4 > *UAS*-dGFPNb:mSc3 | *if/CyO; R23E10-Gal4, UAS-dGFPNb:mSc3/+* |
| Figure S5B-H | See Figure 4 C, D, E | See Figure 4 C, D, E |
| Figure S6C | *RacGAP50C*$^{ALFA\text{-}V5\text{-}GCN4}$ | *RacGAP50C*$^{ALFA\text{-}V5\text{-}GCN4}$ |
| Figure S6D | *RacGAP50C*$^{ALFA\text{-}V5\text{-}GCN4}$, *en*-Gal4>*UAS*-dV5:mSc3 | *RacGAP50C*$^{ALFA\text{-}V5\text{-}GCN4}$*/en-Gal4; dV5:mSc3/+* |
